# A human transcriptomic deletion links cortical expansion and cancer

**DOI:** 10.64898/2026.09.16.752114

**Authors:** Alexander L. Starr, Ana Villalba, Jenna Rever, Yuwei Chen, Florian M. Pauler, Leslie Magtanong, Christopher A. Maxwell, Simon Hippenmeyer, Hunter B. Fraser

## Abstract

Alternative splicing represents a major source of potential evolutionary novelty in protein sequence. Despite this, relatively few studies have investigated the functional impacts of human-specific alternative splicing. Here, we analyze RNA-sequencing data from nine iPSC-derived cell types and identify dozens of human transcriptomic conserved deletions (htCONDELs): evolutionary divergence in splicing that leads to the partial or full removal of conserved protein-coding sequence from the human transcriptome. We investigated one example in detail: an htCONDEL in the gene *HMMR*, which encodes a centrosomal protein that binds microtubules and regulates mitosis. We found that this htCONDEL, which produces a human-specific isoform lacking a conserved microtubule-binding domain, enables cells to flexibly divide with a broader range of mitotic spindle orientations *in vitro*. Introducing the human *HMMR* isoform into mice similarly alters the orientation of cell division in the developing neocortex and increases the production of outer radial glia-like cells, the expansion of which played an essential role in increasing human brain size. Combined with previous work implicating the same *HMMR* isoform in carcinoma progression, our results suggest that evolution of *HMMR* splicing in humans increased plasticity in cell division, potentially leading to tradeoffs between advantageous effects on brain development and deleterious effects on cancer risk later in life.

## Introduction

Alternative splicing, the process by which a single pre-mRNA transcript is processed into multiple distinct mRNA isoforms, affects 95% of human multi-exon genes and vastly increases proteomic diverstiy^1,2^. From an evolutionary perspective, changes in splicing offer a powerful substrate for phenotypic novelty: by altering the inclusion of exonic sequences, a single substitution can have large, potentially cell type-specific effects on protein function^3^. In particular, substitutions that result in the deletion of conserved amino acids might have substantial, phenotypically impactful effects on protein function. While splicing-induced loss of protein sequence in ARHGAP11B has been linked to increased human brain size^4^ and deletions of conserved DNA sequence have been delineated genome-wide^5,6^, there have been limited efforts to systematically identify human-specific alternative splicing that leads to loss of protein sequence, which we refer to as “transcriptomic deletions”. As a result, our understanding of how splicing divergence contributed to uniquely human phenotypes remains limited^7–9^.

Although the transcriptomic deletion of a conserved portion of a protein could generate beneficial phenotypic novelty, such changes are generally expected to be constrained by purifying selection. Therefore, when tolerated or even beneficial in specific developmental or physiological contexts, partial losses of protein sequence may create new vulnerability to disease in other contexts. Such evolutionary tradeoffs are a central prediction of the antagonistic pleiotropy theory, which proposes that alleles favored by natural selection because they improve fitness early in life may have deleterious consequences later in life^10,11^. This framework has been invoked to explain why cancer remains prevalent across most mammals^12^. Advantageous changes in the molecular pathways that promote growth, regeneration, tissue remodeling, and plasticity during development may become deleterious later in life as these same processes are central to tumorigenesis^13–17^. Notably, although epithelial cancers (carcinomas) are common in humans, they appear to be remarkably rare in other great apes, suggesting that humans are more susceptible to carcinoma^18^. This disparity raises the possibility that molecular innovations contributing to uniquely human traits may have simultaneously increased human vulnerability to some types of cancer. However, although a handful of examples have directly linked human-specific sequence changes to both phenotypic innovation and increased disease susceptibility^19,20^, alternative splicing has yet to be implicated in this key aspect of human evolution.

## Main

The generation of evolutionarily novel protein isoforms can occur through two main mechanisms. First, exonic sequence can be gained if new splicing junctions evolve, either leading to a new exon or extending an existing one (Fig. 1a). Second, exonic sequence can be lost if existing splice sites are weakened or a new splice site within an existing exon is strengthened (Fig. 1b). To identify species-specific splicing events, we analyzed published RNA-seq data from nine human-chimpanzee hybrid iPSC-derived cell types^21–23^. These hybrid cells were generated by *in vitro* fusion of human and chimpanzee cells, leading to stable tetraploid cells in which the genomes of both species share the same nucleus^22^. By comparing the splicing patterns of human vs. chimpanzee alleles that are exposed to precisely the same *trans*-acting factors, we isolate the effects of *cis*-acting divergence, i.e. sequence changes that affect splicing of only one species’ alleles. In addition, these hybrid cells control for confounding factors such as differences in cell type composition, developmental stage, or environment^22^. Using this data set, we identified 1,653 species-specific splicing events (at a 5% FDR; see Methods), 1,061 of which have the potential to produce novel proteins (Fig. 1c).

**Figure 1:**
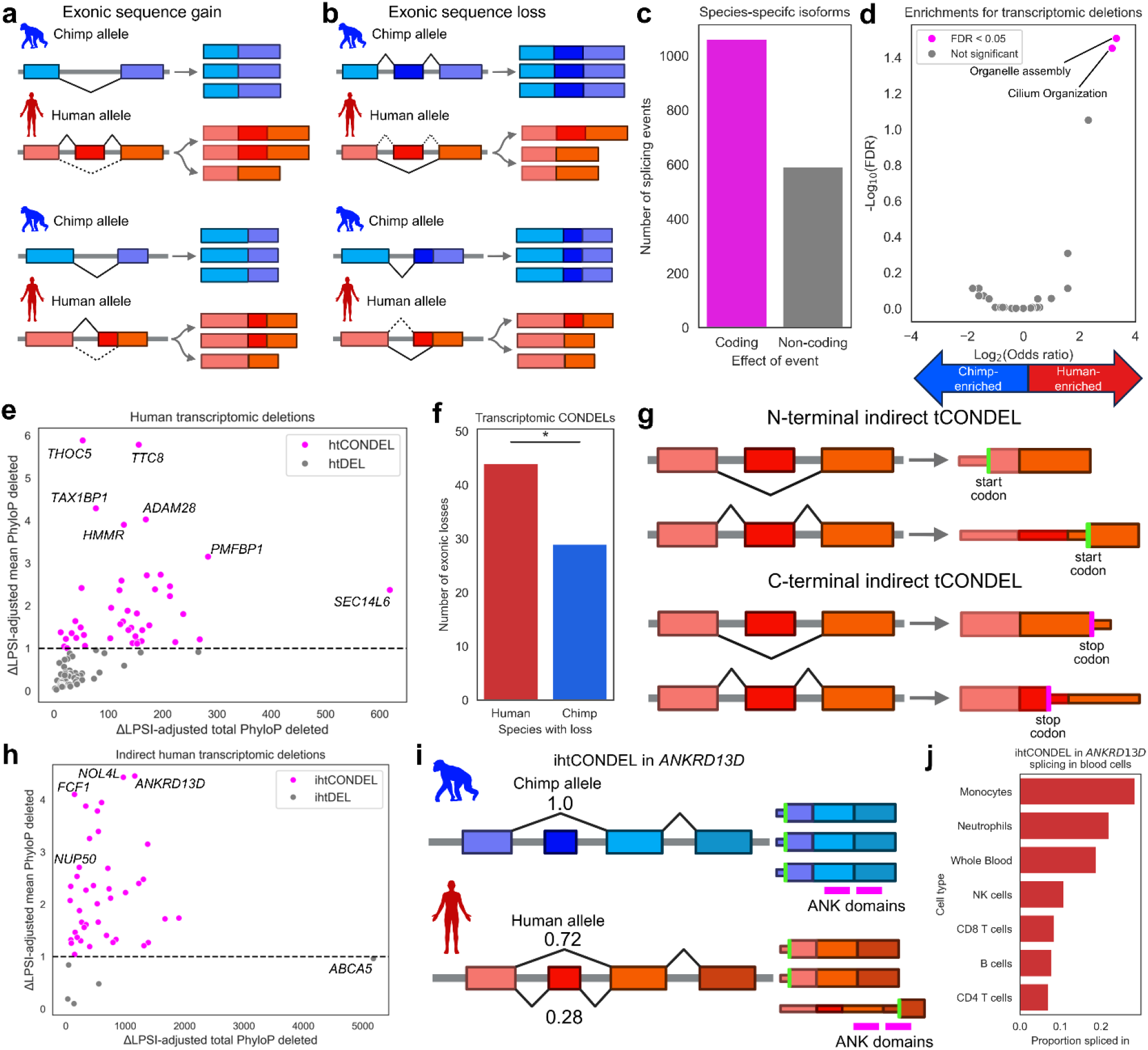
Identification of transcriptomic deletions. **a)** Hypothetical example of exonic sequence gain. Colored boxes represent exons and grey lines represent introns. Black lines indicate splice junction usage, with dashed lines showing the less common splicing usage. Top shows gain of an exon, bottom shows gain of new exonic sequence in a previously existing exon. **b)** Same as in (a) but for exonic sequence loss. **c)** Number of coding and non-coding splicing events that create species-specific isoforms. **d)** Enrichments for human transcriptomic deletions relative to chimpanzee. **e)** Human transcriptomic deletions. Conserved deletions are shown in pink. The x- and y-axes show two different measures of the extent of conserved protein-coding deletion. **f)** Number of human and chimpanzee transcriptomic CONDELs. * = p < 0.05. **g)** Schematic of how indirect tCONDELs may occur. **h)** Human indirect transcriptomic deletions. Indirect conserved deletions are shown in pink. The x- and y-axes show two different measures of the extent of conserved protein-coding deletion. **i)** Schematic of ihtCONDEL for *ANKRD13D*. The gained human splicing leads to the use of a downstream start codon. PSI (proportion spliced in) values are from monocytes. **j)** Proportion spliced in for ihtCONDEL-causing exon inclusion across immune cells for *ANKRD13D*.

To further prioritize likely functional splicing events, we focused on exonic losses, hereafter referred to as transcriptomic deletions (tDELs), that do not disrupt the reading frame. Interestingly, human tDELs (htDELs) are enriched for ciliary functions relative to chimpanzee tDELs (ctDELs) (Fig. 1d). We reasoned that tDELs that remove conserved sequences would be most likely to affect phenotype, similar to the idea underlying human conserved deletions of genomic sequence (hCONDELs^5,6^). Using this strategy, we identified 44 human transcriptomic conserved deletions (htCONDELs) (Table S1, Fig. 1e). In contrast, we identified only 29 chimpanzee tCONDELs (Fig. 1f). Human tDELs were 2.0-fold more likely to be tCONDELs as compared to chimpanzee tDELs (p = 0.015, Fisher’s exact test). For comparison, the few protein-coding human conserved DNA deletions^6^ collectively affect a total of 58 amino acids. In contrast, htCONDELs affect 1061 amino acids. Normalizing by the proportion of transcripts affected (e.g. two amino acids deleted from 50% of transcripts would equal one constitutive amino acid), htCONDELs impact the equivalent of 509 constitutive amino acids. In sum, these analyses suggest that htCONDELs are enriched in the human lineage, and collectively impact a large number of highly conserved amino acids.

### Identification of indirect htCONDELs

Although we are measuring RNA splicing, it is ultimately the effects of splicing on the proportion of protein isoforms that are most likely to affect organismal phenotypes. We reasoned that the exclusion or inclusion of out-of-frame exons could effectively truncate the final protein by leading to the use of an alternative translation initiation site, causing a frameshift, or introducing an early stop codon, in some cases without inducing RNA degradation (Fig. 1g). We term these events indirect htCONDELs (ihtCONDELs). Using a conservative strategy that involves curated human mRNA transcript information (and therefore cannot be applied to identify indirect tCONDELs in chimpanzee; see Methods), we identified 40 ihtCONDELs (Table S2, Fig. 1h). This pipeline guarantees that ihtCONDEL transcripts are supported by curated human reference transcriptomes, largely reducing the potential for identification of spurious transcripts. Moreover, of the 21 ihtCONDEL-linked isoforms that gained protein sequence not found in the canonical proteoform, 13 are experimentally supported by proteomics^24^ and an additional two are supported by ribosome profiling^25^ (i.e. translated, Table S2). This suggests that a sizable fraction of ihtCONDELs produce novel proteins that may have altered functions.

Notably, ihtCONDELs tend to affect many more conserved amino acids than their direct counterparts (Fig. 1e vs. Fig. 1h). For example, *ANKRD13D* is a part of ancient protein family involved in EGF receptor endocytosis that is conserved across metazoans. A human-specific exon in *ANKRD13D* leads to the use of a downstream alternative start codon, removing one and part of another highly conserved ANK domain that determines protein binding specificity (Fig. 1i). Interestingly, the inclusion of this exon is highest in whole blood in adult humans (Fig. S1a) and, among blood cell types, highest in neutrophils and monocytes (Fig. 1j). As these cell types also have the highest *ANKRD13D* expression (Fig. S1b), this suggests that this ihtCONDEL may have affected receptor endocytosis in the immune system during human evolution, a process central to immune evasion by viruses such as cytomegalovirus^26^.

To further prioritize (i)htCONDELs, we scanned the literature for splicing events known to affect protein function or organismal phenotype. Consistent with most alternative isoforms being unstudied, phenotypic effects have been reported for only three ihtCONDELs in *RNMT*^27^, *NVL*^28^, and *NUP50*^29^ and one htCONDEL in *HMMR*^30^. For *NUP50*, the inclusion of a human-specific exon (with the possibility of a chimpanzee-specific exon loss ruled out by this exon also being absent from published macaque datasets; see Methods) results in the removal of 28 highly conserved amino acids (Fig. 2a), producing the protein Npap60S^29^. This truncated version of the protein inhibits release of cargo during nuclear import, whereas the ancestral full length Npap60L promotes release^29^ (Fig. 2a). Strikingly, this exon has moderate inclusion across development in all tissues investigated but is the dominant isoform only in adult testis^31^ (Fig. 2b-c). Although several SNPs are moderately associated with the frequency of this exon’s inclusion (known as splicing QTLs, or sQTLs) in testis^32^, all 410 Genotype-Tissue Expression consortium (GTEx) testes samples, regardless of genotype, include the exon to some extent (Fig. S1c), This suggests that inclusion of this exon at some level is shared by most, or perhaps all, humans.

**Figure 2:**
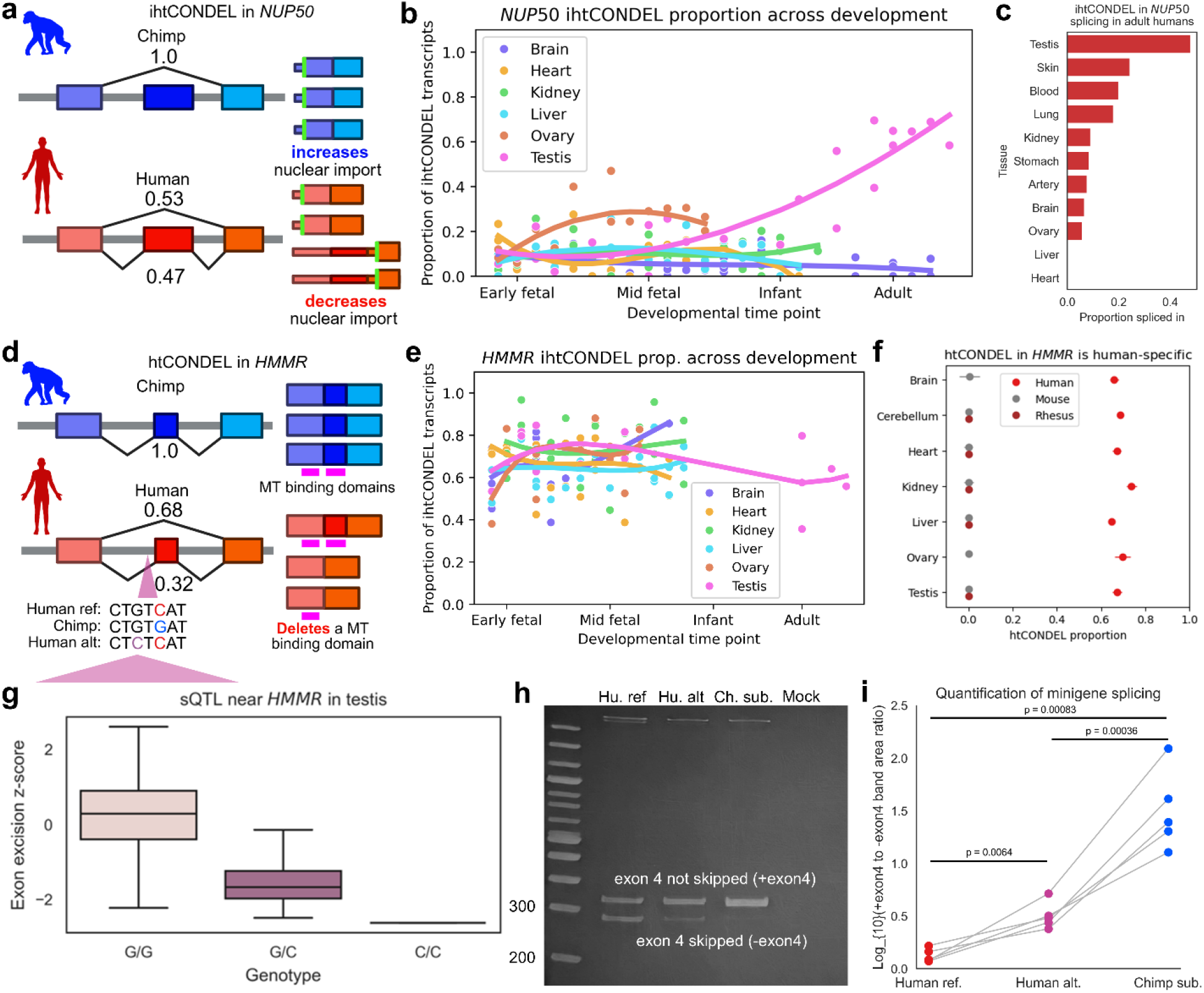
(i)htCONDELs in *HMMR* and *NUP50*. **a)** Schematic of ihtCONDEL in *NUP50*. **b)** Proportion of transcripts that contain ihtCONDEL exon across development in humans. **c)** Proportion of ihtCONDEL exon spliced in across tissues in GTEx. **d)** Schematic of htCONDEL in *HMMR*. The sequences at the bottom show the region of the genome approximately 60 bases upstream of exon 4. **e)** Proportion of transcripts with htCONDEL exon 4 spliced out in *HMMR* across development in humans. **f)** Skipping of *HMMR* exon 4 in mouse and rhesus macaque compared to human. **g)** Exon excision z-score from testes for *HMMR* exon 4 splicing QTL. Negative z-score indicates less exon 4 skipping. **h)** Effects of human sQTL lead SNP alternative allele (pink in (d)) and reversion of human-derived substitution (blue in (d)) to chimpanzee base on exon 4 skipping in *HMMR* in minigene assay. The top band around 300 nucleotides (nt) represents inclusion of exon 4 whereas the lower band around 260 nt represents skipping of exon 4. The faint band at the top is unspliced RNA. **i)** Quantification of effects of substitutions on exon 4 skipping in minigene assay. All values are relative and are not reflective of the actual ratio of isoforms. P-values are from a paired t-test.

For *HMMR*, skipping of exon 4 is known to remove one of two microtubule binding domains^30^ (Fig. 2d). This -exon4 isoform has been implicated broadly in cancer progression^30^ and promotes metastasis of pancreatic ductal adenocarcinoma, whereas the full-length isoform does not^33^. The -exon4 isoform is consistently dominant across human development (Fig. 2e, making up approximately 50-80% of the total *HMMR* transcripts) and human-specific as skipping of exon 4 is observed only very rarely in rhesus macaques and mice across development (Fig. 2f).

To identify the primary causal substitution, we scanned the region within 75 bases of the exon. We identified a human-derived substitution near the branch point (a key region for successful splicing) of exon 4 (Fig. 2d). Remarkably, there is a human polymorphism two bases away that is the lead sQTL for increased inclusion of exon 4 in GTEx^32^ (Fig. 2d,g). Despite the presence of this sQTL, all humans in GTEx skip the exon to some extent (Fig. S1d), again suggesting that this unique splicing event may be shared by all humans. Using a minigene assay, we confirmed that converting the substitution to the ancestral allele and the polymorphism to the minor allele affects splicing of exon 4, with the fixed substitution having a considerably larger effect on exon 4 inclusion (Fig. 2h-i).

### Forced skipping of HMMR exon 4 induces non-planar cell divisions and alters daughter cell phenotypes

To study the effect of exon 4 skipping on the behavior of individual human cells, we used Cas9 to create two distinct genomic deletions in clonogenic, non-malignant, p53-competent MCF10A human mammary epithelial cells expressing endogenous RFP-TUB1AB for microtubule tracking. One was a complete deletion of exon 4 (ΔEX4), and the other deleted 154 bp from intron 3, 10 bp upstream of the lead sQTL (ΔINT3) (Fig. 3a). Redundant clones were isolated (Fig. S2a), and ΔEX4 clones were found to exclusively express the -exon4 isoform (Fig. 3b-c). The full-length *HMMR* isoform comprised 75% of transcripts in parental MCF10A cells, consistent with their heterozygous sQTL genotype (Fig. S1d), versus 23% of transcripts in ΔINT3 cells, and these proportions were also reflected at the protein level (Fig. 3b-c). This suggests that elements in the 154-bp deleted intron 3 region regulate exon 4 splicing.

**Figure 3:**
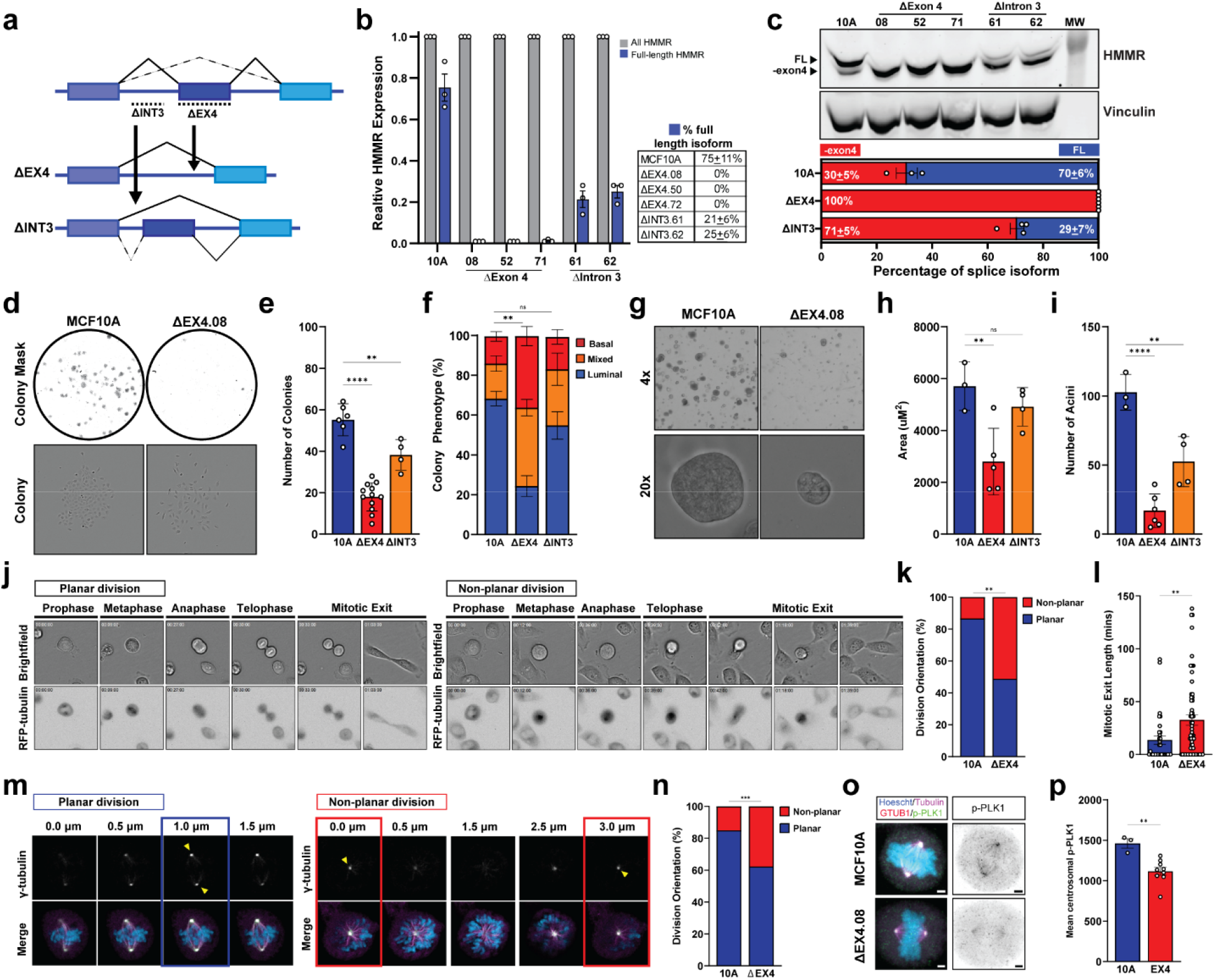
Characterization of effects of *HMMR* exon 4 skipping on cell division *in vitro*. **a)** Diagram depicting the CRISPR editing strategy for ΔEX4 and ΔINT3 clones. **b)** qPCR of *HMMR* transcript isoform abundance, showing full-length isoform specific abundance (blue) relative to the abundance of all HMMR isoforms (grey). Percentage of full length *HMMR* isoform is indicated in the table on the right **c)** Immunoblot for HMMR, with full-length (FL) and -exon4 splice isoform indicated for MCF10A cells and all ΔEX4 and ΔINT3 clones, with Vinculin (VCL) as a loading control. Percentage of full length isoform (blue) and -exon4 isoform (red) are indicated in the bar graph below **d)** Whole well scans of day 5 colony forming assays (above) with representative images of individual colonies (below). **e)** Number of colonies per 100 cells seeded (One-way ANOVA), and **f)** percentage of colony phenotype (Chi-squared test). **g)** Brightfield images of day 9 acini at 4x and 20x objective. **h)** Mean acini area and **i)** number of acini >2000um^2^ per 4x field of view (One-way ANOVA). **j)** Live cell imaging of day 3 MCF10A RFP-TUBA1B colonies using bright-field and RFP (tubulin). Mitotic phase is indicated, showing both planar (left) and non-planar (right) divisions relative to the cell culture substrate. **k)** Percentage of planar and non-planar telophase cells (Chi-squared test, n=20 cells per genotype). **l)** Length of mitotic exit from live cell day 3 MCF10A RFP-TUBA1B colonies (Students t-test n=20 cells per genotype). **m)** Confocal Z-stack images of metaphase cells showing the spindle poles (y-tubulin), and a merged image stained for y-tubulin (white), DNA (blue), and tubulin (purple). Yellow arrows indicate in focus spindle poles, and Z position is indicated above. **n)** Quantification of metaphase spindle angle (Chi-squared test, n=60 cells per genotype). **o)** Confocal immunofluorescence images of metaphase cells stained for DNA (Hoechst), y-tubulin, tubulin, and p-PLK1, with a merged image of all channels, and inverted image of the p-PLK1 channel. **p)** Mean p-PLK1 fluorescence at the metaphase spindle poles (Students t-test). For all plots, statistical tests are indicated in the legend. Statistical significance is represented as follows **** = p<0.0001, *** = p<0.001 ** = p<0.01, * = p<0.05, ns = p>0.05. ΔEX4 datapoints represent mean values for three clones from a minimum of three replicate experiments (n=9) and ΔINT3 datapoints represent mean values for two clones from two replicate experiments (n=4), with one exception: datapoints in panel l represent values from individual mitotic MCF10A and ΔEX4 cells extracted from three replicate experiments.

We then seeded these cells at clonal density to measure their growth phenotypes. Several epithelial tissues are composed of both luminal (i.e. adherent to a substrate) and basal (i.e. away from the substrate) cell subtypes, defined by both phenotypic (Fig. S2b) and molecular profiles^34,35^. Although MCF10A cells typically form phenotypically luminal colonies^36^, ΔEX4 and ΔINT3 clones exhibited reduced colony-forming capacity and were more likely to produce sparse basal colonies with fewer cells after 5 days of adherent growth (Fig. 3d-f, Fig. S2c-d). Similar phenotypic changes were observed for ΔEX4 cells grown under 3D culture conditions (Fig. 3g-i).

The reduced adherent growth phenotype observed in ΔEX4 clones is not a result of reduced cell division, as we observed no differences in the percentage of mitotic cells nor in the distribution of mitotic phases between MCF10A cells and ΔEX4 clones (Fig. S2e-f). Moreover, live-cell imaging (Fig. 3j) revealed no delays in prometaphase, metaphase, anaphase, or telophase (Fig. S2g-h). However, time-lapse imaging of colonies showed ΔEX4 cells divided more frequently at non-planar angles relative to the plate surface, with a subsequent delay in mitotic exit (Fig. 3k-l). Confocal imaging of metaphase spindle pole position confirmed that spindles were oriented at non-planar angles more frequently in ΔEX4 cells (Fig. 3m-n). Collectively, these results indicate that skipping of *HMMR* exon 4 alters spindle positioning to enable non-planar cell division and change clonogenic capacity.

HMMR localizes to spindle microtubules and centrosomes to control the mitotic spindle positioning pathway^37–39^, and exon 4 skipping is known to alter the protein’s localization^40,41^. To gain insight into how *HMMR* exon 4 splicing might alter spindle orientation, we assayed whether skipping of *HMMR* exon 4 alters its localization during cell division or its interacting proteins. We found no difference in mean HMMR fluorescence at metaphase spindle microtubules or mitotic centrosomes between ΔEX4 clones and wildtype MCF10A cells (Fig. S3a-c), consistent with the splicing of exon 4 altering only interphase localization of HMMR^40,41^. Similarly, we found no significant difference in tubulin, y-tubulin, Ac-tubulin, or TPX2 abundance at the spindle (Fig. S3d-i), which are known mitotic interactors of HMMR^42–45^.

However, phosphorylated-PLK1 was significantly reduced at the spindle poles in ΔEX4 cells (Fig. 3o-p, Fig. S3j-k). As chromosomes align during metaphase, centrosome-localized PLK1 locally regulates pulling forces on the mitotic spindle and dictates the fidelity of oriented cell division in a conserved mechanism^46^. Because phosphorylation of PLK1 provides a spindle pole-localized intrinsic code for spindle positioning^46^, it is plausible that forced skipping of HMMR exon 4 mechanistically reduces this intrinsic signal with consequent augmentation of non-planar cell division and altered daughter cell phenotypes in vitro. Importantly, non-planar (asymmetric) cell division is also an essential driver of tissue homeostasis and differentiation in many tissues, especially in brain^47^.

### Humanization of *Hmmr* exon 4 affects spindle orientation in the developing mouse brain

Given that both spindle orientation^48,49^ and HMMR^37,50^ play an essential role in the development of the neocortex, we wanted to evaluate whether skipping of exon 4 might have affected human brain evolution. To do this, we generated a humanized exon 4 mouse model in which exon 4 and flanking intronic sequence was replaced with the human reference sequence (Fig. 4a, two lines from independent founders were analyzed). Throughout, we used the term wildtype to refer to *Hmmr*^+/+^ mice (i.e. mice that have the canonical mouse sequence for exon 4 and the flanking intronic regions) and humanized to refer to the *Hmmr*^HEx^^4/HEx^^4^ mice which have exon 4 and the flanking human intronic regions humanized. Consistent with our minigene results, we found that humanized mice skipped exon 4 in 60-61% of transcripts compared to less than 2% in wildtype mice (Fig. S4a).

**Figure 4:**
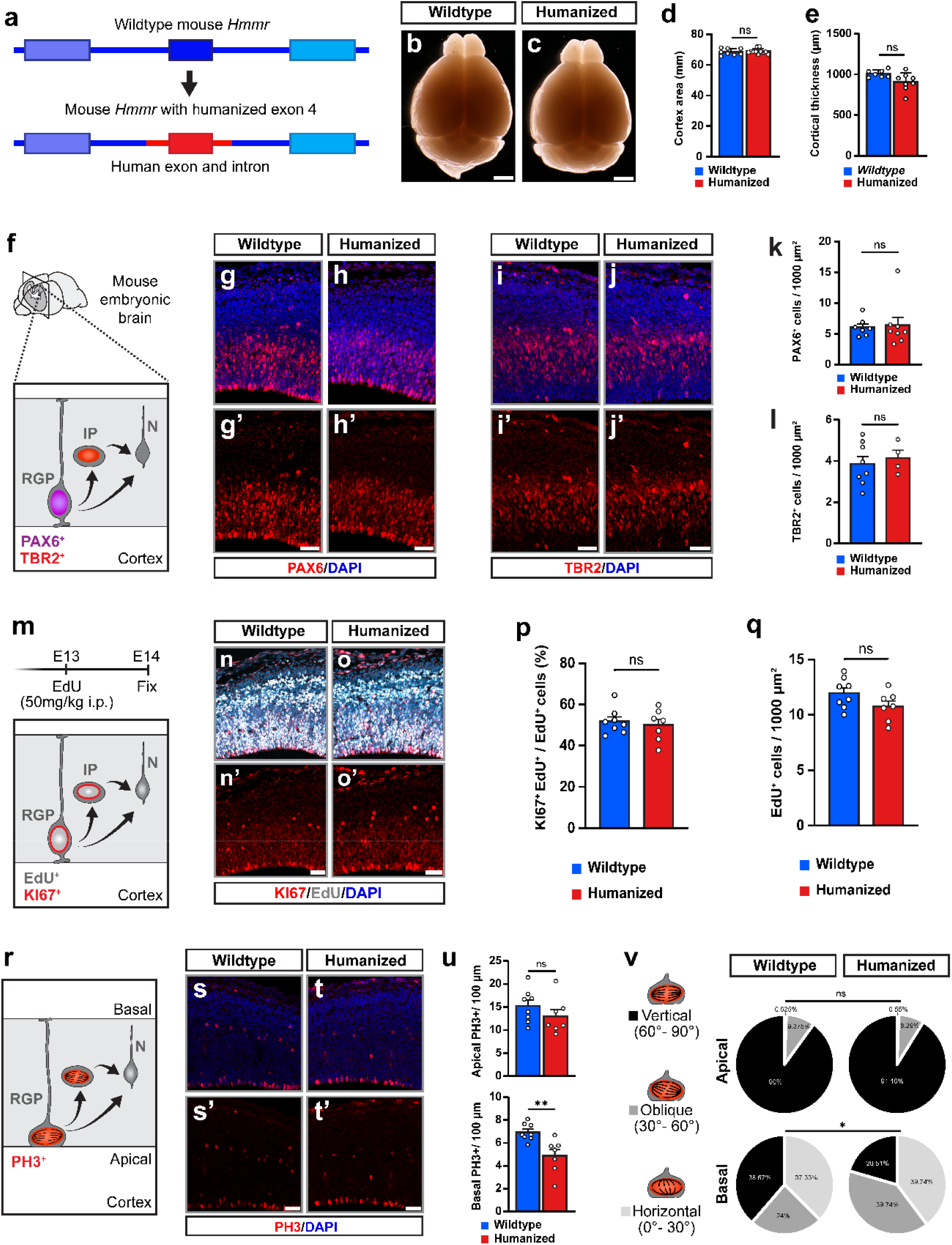
Humanizing Exon 4 shifts basal progenitors spindle orientation without affecting brain size or apical RGP proliferation. **a)** Schematic of the generation of mice carrying a human *Hmmr* exon 4. The dark blue box indicates mouse exon 4, whereas the red insert corresponds to the human exon 4 sequence and flanking intronic regions. **b)** *Hmmr* wildtype P21 mouse brain image. **c)** *Hmmr* carrying humanized exon 4 P21 mouse brain image. **d)** Quantification of the cortical area measured from stereomicroscope images (µm^2^). **e)** Quantification of cortical thickness in µm. **f)** Schematic of progenitor proliferation in the embryonic mouse cortex. Radial glial progenitors (RGPs; Pax6⁺) residing in the ventricular zone divide to expand the progenitor pool and give rise to intermediate progenitors (IPs; Tbr2⁺) and neurons (N), contributing to cortical growth. **g-h)** Representative images of *Hmmr* wildtype and humanized mouse somatosensory cortex, DAPI (blue) and PAX6 (red). **i-j)** Representative images of *Hmmr* wildtype and humanized mouse somatosensory cortex, DAPI (blue) and TBR2 (red). **k)** Quantification of PAX6^+^ cells per 1000 µm^2^ in the somatosensory cortex of E14 wildtype and mutant embryos. **l)** Quantification of TBR2^+^ cells per 1000 µm^2^ in the somatosensory cortex of E14 wildtype and mutant embryos. **m)** Illustration of the cell cycle exit assay. Pregnant females were intraperitoneally injected with EdU (50 mg/kg) at E13, and embryos were collected 24 h later at E14. During this period, EdU-labeled radial glial progenitors (RGPs) generate intermediate progenitors (IPs) and neurons (N), which inherit the EdU label. KI67 immunostaining identifies cycling cells, including RGPs and IPs, allowing the distinction between proliferating (EdU⁺KI67⁺) and cell cycle–exited (EdU⁺KI67⁻) cells. **n-o)** Representative images of *Hmmr* wildtype and humanized mouse somatosensory cortex, DAPI (blue), EdU (grey) and KI67 (red). **p)** Quantification of the cell cycle re-entry (KI67^+^EdU^+^ cells per total number of EdU^+^ cells) in the somatosensory cortex of E14 wildtype and mutant embryos. **q)** Quantification of the total number of EdU^+^ cells per 1000 µm^2^ in the somatosensory cortex of E14 wildtype and mutant embryos. **r)** Schematic of G2/M-phase labeling using phospho-histone H3 (PH3). Apical PH3⁺ cells at the ventricular surface correspond to mitotic radial glial progenitors (RGPs), whereas basally located PH3⁺ cells represent dividing secondary progenitors. **s-t)** Representative images of *Hmmr* wildtype and humanized mouse somatosensory cortex, DAPI (blue) and PH3 (red). **u)** Quantification of the number of apical or basal PH3^+^ cells per 100 µm of ventricular surface in the somatosensory cortex of E14 wildtype and mutant embryos. **v)** Quantification of spindle orientation in apical and basal PH3^+^ mitoses. Spindle angles were categorized as vertical (60°–90°), oblique (30°–60°), or horizontal (0°–30°) relative to the ventricular surface. While spindle orientation in apical progenitors was unchanged between genotypes, humanized basal progenitors exhibited a statistically significant shift toward oblique and horizontal spindle orientations compared with wildtype controls (p = 0.027). Bars and error bars represent mean ± SEM. Statistical significance was assessed using unpaired t test (d, e, k, I, p, q, u) or Chi-square test (v). Scale bars, 2mm (b, c) or 40µm (g, h, l, p). Nuclei were counterstained using DAPI (blue).

Because previous work has shown that deletion or truncation of *Hmmr* causes gross differences in brain morphology^37,50^, we first investigated morphology in adult brains. We observed no clear differences in brain size, cortical area, or cortical thickness between humanized and wildtype mice (Fig. 4b-e). Furthermore, we found no differences in the density of radial glia (marked by PAX6)^51^ or intermediate progenitors (marked by TBR2)^51^ (Fig. 4f-l). By injecting EdU 24 hours prior to fixation we investigated cell cycle exit at E14, using EdU and KI67 labeling to mark recently proliferating cells (EdU+) and calculating the fraction of those that remain cycling (KI67+ and EdU+). We found no difference in the ratio of KI67+ EdU+ (i.e. mitotic EdU+ cells) to EdU+ cells nor in the density of EdU+ cells (Fig. 4m-q), suggesting that, as in the MCF10A cells, there are minimal alterations to cell proliferation when exon 4 is skipped. Collectively, this suggests that the naturally occurring change in *Hmmr* splicing we have identified has more subtle effects than artificial deletions of either the full gene or larger portions of the protein^37,50^.

As the primary effects of exon 4 skipping *in vitro* were on actively dividing cells, we next explored whether mitosis was altered in humanized mice. While humanized mice had no difference in the density of actively dividing apical progenitors (i.e. PH3+ cells), they had significantly fewer basal divisions (Fig. 4r-u). Strikingly, the humanized mice also had a significantly higher proportion of basal progenitors dividing at oblique or vertical angles relative to wildtype mice, mirroring the differences in the orientation of cell division observed *in vitro* (Fig. 4v). In contrast, there was no difference in the spindle plane of apical progenitors (Fig. 4v), suggesting that the effects of exon 4 skipping are specific to basal progenitors.

### Humanized *Hmmr* exon 4 mice have a larger number of outer radial glia-like cells

To determine whether humanization affects cell population composition and/or induces molecular changes, we generated single cell RNA-sequencing data (scRNA-seq) from embryonic day 14 mouse forebrain and bulk RNA-seq data from purified populations of progenitors (Fig. 5a-b) from wildtype and humanized mice. In the scRNA-seq data, we identified all expected cell types (Fig. 5c, Fig. S4b) and focused analysis on radial glia (marked by *Pax6*, Fig. 5d). Many radial glia express *Hmmr*, as expected given its broad expression in dividing cells^52^ (Fig. 5e). Next, we tested for differential expression between humanized and wildtype mice in radial glia and the bulk progenitor RNA-seq data. In both cases, expression was quite similar: there were no genes with individually significant differential expression (all FDRs > 0.05), suggesting any transcriptional effects of exon 4 skipping are subtle. However, we observed considerable agreement in differential expression between the bulk and scRNA-seq data (which are from different mice from independent litters), suggesting that the subtle transcriptional effects of *Hmmr* exon 4 skipping in the developing brain are nevertheless reproducible (Spearman’s rho = 0.49, p = 0.0019, Fig. 5f). We focused further analysis on genes that had the same sign of log_2_ fold-change in the bulk and scRNA-seq data. For example, HOPX—a canonical marker of oRG—was slightly more highly expressed in both of our humanized RNA-seq data sets, so it was included for further analysis even though it did not reach statistical significance in either data set individually.

**Figure 5:**
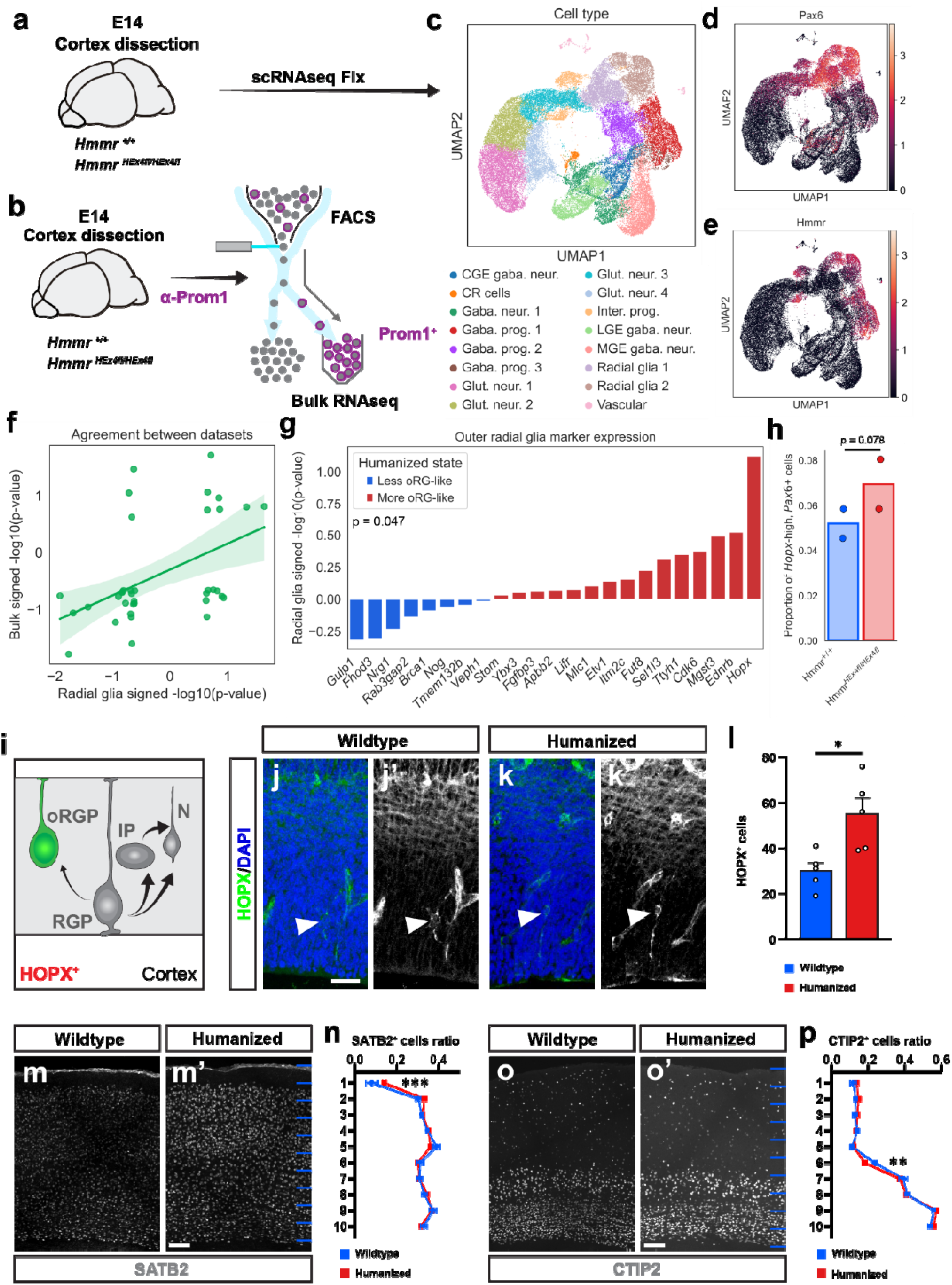
Skipping of *Hmmr* exon 4 increases the abundance of outer radial glia-like cells (oRGPs). **a)** Experimental strategy to generate single cell suspension from wildtype and mutant dissected cortices at embryonic time point E14 followed by scRNA-seq Fix protocol. **b)** Experimental strategy to FACS-enrich for cortical apical RGPs (Prominin^+^). Prom^+^ cells were isolated from wildtype and mutant dissected cortices at embryonic time point E14 for Bulk RNA-seq assessment. **c)** UMAP of cell types identified in E14 scRNA-seq data. C/L/MGE = caudal/lateral/medial ganglionic eminence, CR = Cajal-Retzius, neur. = neurons, prog. = progenitors, Glut. = glutamatergic, Inter. = intermediate. **d)** Expression of *Pax6* on UMAP. **e)** Expression of *Hmmr* on UMAP. **f)** Agreement between bulk and scRNA-seq differential expression quantified using the -log_10_(p-value) multiplied by the sign of the log_2_ fold-change between wildtype and humanized mice. Line shows result of linear regression and 95% confidence interval. **g)** Differential expression of outer radial glia progenitor (oRGP) marker genes from scRNA-seq data. Red indicates higher expression in humanized mice. **h)** Proportion of radial glia that are *Hopx*-high. Dots represent samples and p-value is from paired t-test with pairing based on the initial founder humanized mouse used. **i)** Schematics showing that, although in very low numbers, mouse RGPs also generate another type of secondary progenitor typically more abundant in human cortex (oRGPs; HOPX^+^). **j-k)** Representative images of HOPX^+^ cells in *Hmmr* wildtype and humanized somatosensory cortex, DAPI (blue) and HOPX (green/grey). White arrows point to HOPX^+^ cells. **l)** Quantification of the number of HOPX^+^ cells in 5 cortical sections of E14 humanized and wildtype embryos. **m)** Representative images of SATB2^+^ cells (grey) in P21 *Hmmr* wildtype and humanized somatosensory cortex. **n)** Quantification of SATB2^+^ cells distribution along 10 bins in P21 wildtype and mutant somatosensory cortex. **o)** Representative images of CTIP2^+^ cells (grey) in P21 *Hmmr* wildtype and humanized somatosensory cortex. **p)** Quantification of CTIP2^+^ cells distribution along 10 bins in P21 wildtype and mutant somatosensory cortex. Bars and error bars represent mean ± SEM. Statistical significance was determined by using unpaired t test (HOPX) or two-way ANOVA (SATB2 and CTIP2). Scale bars, 40µm (HOPX) and 100µm (SATB2 and CTIP2). Nuclei were stained using DAPI (blue).

In mammals, basal progenitors consist of at least two distinct populations: TBR2+ intermediate progenitors that typically (though not exclusively) undergo a single cell division that produces two neurons^53,54^, and HOPX+ basal or outer radial glia progenitors (oRGP) that can self-renew to produce additional progenitors as well as neurons^55,56^. While oRGP-like cells are very rare in mice, they are common in larger-brained mammals^57,58^. In particular, the number of oRGP is thought to be considerably higher in the developing human neocortex relative to chimpanzees and other apes^59,60^. While the human-specific gene *ARHGAP11B* likely contributed to increased human brain size by increasing the number of outer radial glia^4^, our understanding of how the pool of oRGP may have expanded in humans remains incomplete.

We analyzed these data to evaluate whether humanization of *Hmmr* exon 4 might have increased the number of oRGP-like cells in two ways. First, we found that humanized mouse progenitors expressed significantly higher levels of oRGP markers from humans and mice (Fig. 5g, p = 0.047 comparing the signed differential expression p-value for oRGP markers to all other genes using the Mann Whitney U Test). Second, in the scRNA-seq data, we found that there were more *Hopx*-high, *Pax6*+ (a combination frequently used to identify oRG^55^) in the humanized mice, although this did not reach statistical significance, potentially due to low sample size (Fig. 5h, p = 0.078, one-sided paired t-test). To test this histologically, we stained for HOPX and PAX6 in E14 mouse brain slices. Strikingly, we found that there were significantly (1.8-fold) more HOPX+ PAX6+ cells in humanized than wildtype mice (Fig. 5i-l).

We next tested whether there might still be detectable consequences of this in the adult mouse brain, despite the rarity of oRGP-like cells in mice. Previous work has linked oRGP expansion to disproportionate expansion of upper layer neurons (marked by SATB2) relative to the number of deep layer neurons (marked by CTIP2)^58^. Consistent with this, we found a significantly increased relative number of SATB2+ neurons in the superficial cortical areas of the humanized mice (Fig. 5m-n), accompanied by a significant decrease in CTIP2+ neurons in lower cortical layers (Fig. 5o-p). Collectively, these results show that humanization of exon 4 skipping specifically expands the pool of HOPX+ progenitors in the developing mouse brain.

Our results establish (i)htCONDELs as a distinct and underexplored class of human molecular divergence in which partial loss of protein-coding sequence due to changes in RNA splicing, rather than genomic DNA deletion, drives phenotypic divergence. We validate that one htCONDEL in *HMMR* alters the plane of cell division in (neuro)epithelial cells and increases the abundance of outer radial glia-like cells in a mouse model. Notably, prior work has shown that the isoform of *HMMR* established as human-specific in this work contributes to the progression of multiple cancers^33,61–66^, particularly pancreatic ductal adenocarcinoma. Based on these observations, the htCONDEL we identified in *HMMR* may have contributed to the apparent increased susceptibility of humans to carcinoma relative to other great apes^18^.

These results suggest that human-specific molecular changes can generate evolutionary tradeoffs by relaxing constraints on a core cellular process such as cell division^67^. In the case of *HMMR*, increased flexibility in spindle orientation may have enabled novel developmental trajectories by increasing the range of cellular behaviors available during neurogenesis. This might also have consequences in tissues where division orientation is critical for maintaining architecture, including the skin^68,69^, intestine^70^, kidney^71^, mammary gland^72^, lung^73^, and testis^74^. At the same time, the human-specific *HMMR* isoform likely increased susceptibility to cancer, which is often characterized by loss of tissue architecture and high flexibility in cellular behavior^15–17^. In this context, altered spindle orientation could allow cancerous or pre-malignant cells to more easily escape architectural constraints and tolerate errors in cell division, ultimately promoting disorganized growth and dissemination into new microenvironments. It is worth noting that spindle orientation is particularly important for carcinoma progression as the loss of tissue architecture caused by aberrant spindle orientation is thought to be an important driver of carcinogenesis and the epithelial to mesenchymal transition^75^. The humanized *Hmmr* mice generated in this study enable testing of these hypotheses and provide a new tool to better model human cancer progression in mice. Overall, our results suggest that evolutionary changes that permit greater developmental and cellular plasticity may create both opportunities and liabilities. If the developmental consequences of *HMMR* exon 4 skipping were adaptive— such as its potential contribution to human cortical expansion—this htCONDEL would represent a rare example of the antagonistic pleiotropy hypothesized to underlie increased late-life disease risk in humans^10,11^.

More broadly, *HMMR* represents only one of dozens of (i)htCONDELs identified in this study, several of which remove larger or more evolutionarily conserved portions of proteins in a greater proportion of transcripts. These findings suggest that splicing-mediated deletion of protein sequence represents a previously underappreciated mechanism through which human-specific phenotypes can evolve. We anticipate that future functional characterization of additional (i)htCONDELs will provide further insight into the molecular basis of human-specific traits and evolutionary tradeoffs with disease susceptibility.

## Methods

### Quantification of alternative splicing in cells derived from human-chimpanzee hybrid iPSCs

We used our previously published pipeline^23^ to re-align and process all RNA-seq data from human-chimpanzee hybrid iPSC-derived cells to the human hg38^76^ and chimpanzee PanTro6^77^ reference genomes. Briefly, we used STAR v2.7.8a^78^ to align the data to both the human and chimpanzee reference genomes separately, picard to remove duplicate reads, and Hornet (a modified version of the software package WASP^79^ that removes read overlapping indels) to correct for mapping bias. We then split reads into those from the human and chimpanzee alleles based on overlap with substitutions inferred from the human-chimpanzee genome-wide alignment and filtered using whole genome sequencing data from the parental iPSC lines to only include substitutions homozygous for the respective reference alleles in all lines^23^. We analyzed data from nine different cell types/organoids: induced pluripotent stem cells (iPSC), hepatic progenitors (HP), pancreatic progenitors (PP), skeletal myocytes (SKM), cardiomyocytes (CM), motor neurons (MN), retinal pigmented epithelial cells (RPE)^23^, day 100 cortical organoids (CO)^22^, and cranial neural crest cells (CNCC)^21^.

To quantify splicing, we used the leafcutter^80^ pipeline. We used regtools^81^ v0.5.2 with parameters -s 1 -a 8 -m 50 -M 500000 to quantify read counts overlapping splice junctions for the human and chimpanzee alleles separately. We then used the script leafcutter_cluster_regtools.py to create clusters of overlapping splice junctions across all samples within each cell type. We then used leafcutter_ds.R to test for differential splicing between the human and chimpanzee alleles with the arguments -e human_all_exons.txt.gz --num_threads 4 -g 3 -i 3 -c 5. We only used the -e argument for human-referenced data and human_all_exons.txt.gz represented all Gencode human exons. To visualize the results for the human-referenced data, we used leafviz and prepared data by running prepare_results.R -f 1 for all cell types.

To eliminate false differential splicing calls caused by mapping bias, we lifted over (using LiftOver^82^) each junction from the human-referenced data to the chimpanzee genome and back to the human genome and the chimpanzee-referenced data to the human genome. We then used bedtools^83^ to intersect the lifted-over splice junctions together. For the next step, we retained any cluster of splice junctions for which all junctions successfully reciprocally lifted and each junction from the human-referenced data had a matching initially chimpanzee-referenced junction within 2 base pairs. Next, we removed all junctions with leafcutter proportion spliced in (LPSI, which is distinct from how PSI is typically computed for a single exon as it refers to intron excision rates rather than exon inclusion rates) less than 0.05 for both the human and chimpanzee alleles. We then iterated through all clusters of junctions and removed those for which the difference in ΔLPSI (i.e. the difference in LPSI between the human and chimpanzee alleles) for the human-referenced data and the chimpanzee-referenced data exceeded 0.1 as these were likely the result of mapping bias rather than *bona fide* differences between the alleles. Finally, for all remaining clusters, we normalized LPSI for each allele to 1 (respectively) and recomputed ΔLPSI. We used the resulting tables in the identification of (i)htCONDELs. Hereafter, we only used the values computed for the human-reference data.

### Identification and analysis of htCONDELs

It is worth noting that, unlike DNA deletions, it is difficult to systematically polarize transcriptomic deletions (tDELs) to the human or chimpanzee lineage due to the lack of suitably deep RNA-sequencing data for comparison to gorillas or another great ape. Therefore, we define transcriptomic deletions relative to the other species. As a first, we classified splicing events involving 2, 3, or 4 junctions as either alternative splice site usage (Alt SS), exon skipping, multiple Alt SS (i.e. when there were 3 junctions and it did not match exon skipping), mutually exclusive exon usage, and exon skipping with an Alt SS. We excluded mutually exclusive exon usage as well as splicing events with greater than 4 junctions. From these, we selected a representative intron (i.e. length of DNA between two splice junctions). For Alt SS or multiple Alt SS, this was the longest intron. For exon skipping or exon skipping with an Alt SS, it was the intron that included the exon being skipped. We then considered any splicing event for which the cluster had an FDR < 0.05 and where the representative intron had LPSI for one allele greater than 0.25 and LPSI for the other allele less than 0.025 or had LPSI greater than 0.975 for one allele and less than 0.75 for the other, effectively retaining only splicing events that lead to species-specific isoforms.

From this set of species-specific isoforms, we considered a splicing even to be a human tDEL (htDEL) if it resulted in decreased inclusion of exonic sequence in humans relative to chimpanzees and vice versa for chimp tDELs (ctDELs). This effectively creates two criteria by which something is classified as a tDEL, which we illustrate with htDELs. In the first, exonic sequence that is never spliced out in chimpanzees is frequently spliced out in humans (*HMMR* is an example of this). In the second, exons that are sometimes spliced out in chimpanzees are constitutively spliced out in humans (*TAX1BP1* is an example of this, see Table S1). To identify conserved tDELs (i.e. tCONDELs), we first intersected the representative introns with human Gencode CDS annotations and retained only those that lie between two protein coding exons (they need not intersect a protein-coding exon directly as this would miss cases in which exons constitutively skipped in human are not annotated in Gencode). Next, we identified the region of the genome that is spliced out in htDELs and ctDELs and intersected this with our previously computed PhyloP^84^ scores^85^, a measure of evolutionary conservation for which higher values indicates greater conservation and negative values indicates more rapid evolution than expected by chance. We rank tDELs by two criteria. The first is the sum of the non-negative PhyloP scores for a region, multiplied by the ΔLPSI (sum metric). This effectively weights by the amount of conserved sequence that is spliced out. The second is the mean non-negative PhyloP score for a region, multiplied by the ΔLPSI (mean metric). This instead weights by the average conservation of the sequence spliced out. Ultimately, we consider anything for which the latter metric is greater than 1 to be a conserved deletion (CONDEL). For Alt SS, we additionally required that the splicing event affect more than 6 bases of exonic sequence.

Tissue-level PSI estimates for each cassette exon were derived from the GTEx^32^ v10 median junction read counts (https://gtexportal.org/api/v2/expression/medianJunctionExpression). For each gene, three junctions were queried: the upstream inclusion junction (J_e_p__), the downstream inclusion junction (J_e_n__), and the exon-skipping junction (J_sk_). PSI was computed from median counts as: PSI = (J_e_p__ + J_e_n__) / (J_e_p__ + J_e_n__ + 2 × J_sk_) (the standard formula for PSI^86^, which is distinct from the intron-excision-style PSI from leafcutter^80^). Per-sample PSI and gene expression were computed for *ANKRD13D* across sorted human blood cell types profiled in GSE74246^87^. Junction count matrices were downloaded from recount3^88^ and per-sample PSI was computed using the formula above. Samples with a cluster total (J_e_p__ + J_e_n__ + J_sk_) below 10 reads were excluded as insufficiently covered. Per-sample PSI for the *ANKRD13D* cassette exon was computed across seven sorted human blood cell types profiled in GSE60424^89^ using the same strategy. Gene-level expression was quantified from the recount3 gene sums file and converted to reads per kilobase per million (RPKM). Per-sample normalized excision ratios (NER) and sQTL statistics were retrieved from the GTEx v10 sQTLs^32^ (https://gtexportal.org/api/v2/association/dynsqtl).

For *HMMR*, the sQTL was for the variant chr5_163467648_G_C_b38 and junction chr5:163464802:163469641:clu_98294_+ from testis whereas for *NUP50* this was for chr22_45171031_C_A_b38 and chr22:45168246:45171600:clu_45031_+ junction from lower leg skin. To plot the PSI distribution for the exons in *HMMR* and *NUP50*, we downloaded the full set of junction counts from GTEx v10^32^, extracted the counts for the relevant junctions, computed PSI per individual as described above, and then plotted the histogram. To plot PSI over development in humans and aggregated across development per tissue (the latter for *HMMR* only), we lifted the coordinates of the exon over^82^ from hg38 to hg37, downloaded the human, macaque, and mouse splicing data^31^ from https://apps.kaessmannlab.org/alternative-splicing/, restricted to the exon of interest, and plotted PSI. We additionally used this data to verify that the skipping of *HMMR* exon 4 and the exon that causes the ihtCONDEL in *ANKRD13D* were human-specific by identifying the exon in the Mazin et al. data and finding either that it was almost never skipped in all other species (*HMMR*) or that it did not exist in any other species (*ANKRD13D*). To test for tDEL enrichments between human and chimpanzee, we intersected all Gene Ontology biological process categories^90^ with the list of htDEL genes and ctDEL gene and used Fisher’s exact test.

To test whether the exon in *NUP50* was human-specific, we identified the orthologous genomic region in macaque and downloaded raw fastq RNA-seq data from diverse macaque tissues^91^ (NHPRTR PRJNA261940) and positive control human data^92^ (SRR1551047). We designed unique (i.e. present only once in the genome and transcriptome) 25mers^93^ overlapping either the 190 base pair putatively human-specific exon in *NUP50* or the exon-exon junction of the two flanking exons putatively present in both species. We then grepped for these kmers from the raw fastq files and removed any reads that overlapped intronic sequence. For macaque, we observed 0 reads that included the 190 base pair exon compared to 1,769 that included the exon-exon junction resulting from skipping the 190 base pair exon. In contrast, in humans we identified 40 reads with evidence for inclusion of the 190 base pair exon and 25 with evidence for exclusion, confirming that the methodology used could detect alternative splicing of this exon.

### Identification and analysis of ihtCONDELs

To identify indirect htCONDELs (ihtCONDELs) we first filtered to all species-specific isoforms (see above) that were between two coding exons or intersected a coding exon in the human Gencode annotation. We then further restricted to splicing events for which the region spliced out/in was fully contained in an exon of a protein-coding Gencode v45^94^ or RefSeq^95^ GRCh38.p14 transcript (restricting to NM_ accessions for the latter) and the CDS for at least one of the transcripts containing that exon was shorter than the CDS for the canonical transcript. If multiple transcripts are supported in this manner, we selected the transcript with the shortest CDS. The canonical transcript was selected according to the following criteria, with the search stopping after success: (1) the MANE Select transcript (the search stopped here for > 99% of protein-coding genes), (2) the APPRIS principal 1 transcript, (3) a transcript with TSL 1 and CCDS, or (4) the longest protein-coding transcript.

We then removed all events that had already been classified as an htDEL or ctDEL, restricted to transcripts with either annotated alternative start codons, annotated alternative stop codons, or in which the htDEL affected the first/last exon or skipped the second to last exon (which would not induce nonsense-mediated decay [NMD] as it would lead to a frameshift in the last protein-coding exon). This ensured that the production of an alternative protein product, rather than NMD, was the likely outcome of the splicing event. It is this key quality control step that prevents us from attempting to identify ictCONDELs as this relies on the highly curated reference transcriptomes available for humans but not chimpanzees. We further removed splicing events that required skipping of multiple exons to maintain the reading frame and did not result in a frameshift or truncation more broadly (based on the gtf annotations).

For the events meeting these criteria, we then extracted the coordinates of all CDS exonic regions of the canonical transcript that are not annotated as CDS in the truncated isoform and must not be CDS in order to avoid nonsense-mediated decay (this criterion removed, for example, downstream exons that happen not to be included in the truncated isoform but not as a direct result of the ihtCONDEL), deduplicating all regions assigned to each individual splicing event to prevent double counting lost CDS sequence. We then intersected this bed file with the bed file of PhyloP scores and computed the sum and mean metrics as described above and considered any qualifying event with a mean metric greater than one to be an ihtCONDEL and all other such events to be ihtDELs. To identify splicing effects with known effects, we used an LLM to search for functionally annotated splicing events in each gene, attempted to match these events to the exonic regions spliced out in our (i)htCONDELs, and then retained the four (*HMMR*, *NUP50*, *RNMT*, and *NVL*) with exact matches.

### Validation of ihtCONDELs with proteomics and ribosome profiling

Validating ihtCONDELs required identifying what novel amino acid or translated sequence appeared in the ihtCONDEL transcript (or resulting protein) but not in the canonical transcript so that these novel peptides and/or translated sequence could be scanned for in proteomics or ribosome profiling datasets respectively. For each event, the canonical transcript was identified using the MANE Select mapping (GENCODE v50 / NCBI RefSeq). When MANE Select was unavailable, the Ensembl canonical transcript or the longest CDS protein-coding transcript was used. The transcript ID of ihtCONDELs was chosen as the GENCODE transcript that included the ihtCONDEL splicing event and the fewest splicing differences from the canonical transcript.

If instead the ihtCONDEL transcripts only appeared in RefSeq, the same criteria were used to select the RefSeq transcript. To assign Uniprot IDs to the proteoforms resulting from ihtCONDELs, we manually searched the Uniprot entry for each gene and determined whether the difference in protein sequence resulting from each ihtCONDELs was an annotated Uniprot proteoform. Novel amino acid sequences were derived from the CDS set difference between the novel and canonical transcripts. CDS exons for both transcripts were extracted from GENCODE v50 (or UCSC refGene for RefSeq transcripts). The genomic coordinate set difference (novel CDS minus canonical CDS) yielded the novel-specific coding sequence, which was translated using the standard codon table, with the stop codon excluded from novel region. All results from this pipeline were verified manually in the UCSC genome browser.

For proteomics validation, we performed *in silico* trypsin digestion of the novel amino acid sequences (with up to 2 missed cleavages). We then searched the resulting peptides on the ProteomicsDB web portal^24^. If a peptide exactly matching a novel subsequence appeared in the database, we considered that to be proteomically validated. Any remaining ihtCONDELs with novel amino acid sequence were considered proteomically unvalidated. For ribosome profiling– based validation, we selected ihtCONDELs satisfying two criteria: (1) a maximum per-base ribosome profiling coverage of ≥10 reads within the ihtCONDEL and (2) a ratio of ribosome profiling coverage of the novel translated sequence to that of the nearest canonical exon of >0.1. The first criterion was used to ensure sufficient sequencing coverage to detect translation of the novel sequence, whereas the second required the novel sequence to exhibit appreciable translation relative to the corresponding canonical isoform. Mean ribo-seq coverage was computed over all positions in each the novel and canonical regions, treating positions with no data (NaN/None) as zero.

### *HMMR* exon 4 minigene assay design

We synthesized the hg38 sequence for *HMMR* exon 3, intron 3, exon 4, intron 4, and exon 5 (Genscript) and cloned it into a plasmid backbone, driven by the CMV promoter. We used site directed mutagenesis (Genscript) to introduce two point mutations, chr5:163467650-C-G (reversion to the ancestral allele at the human-derived, fixed substitution) and chr5:163467648-G-C (the alt allele of the sQTL). We refer to these minigenes as reference, ancestral, and sQTL alt respectively. We verified all plasmids with long read sequencing before use (Plasmidosaurus).

### Cell culture for minigene assay

NIH/3T3 cells (cat no. CRL-1658) were obtained from American Type Culture Collection (Manassas, VA). Cell lines were tested for mycoplasma using MycoStrip (cat no. rep-mys-10, InvivoGen, San Diego, CA) and tested negative. NIH/3T3 cells were cultured in Dulbecco’s Modified Eagle Medium (DMEM, cat no.10569010, Thermo Fisher, Waltham, MA), 10% fetal bovine serum (FBS, cat no. 16000044, Thermo Fisher), and 0.5 U/mL penicillin-streptomycin (1x P/S; cat no. 15070-063, Thermo Fisher). Trypsin (cat no. 5200072) was from Thermo Fisher. Phosphate buffered saline (cat no. P5368) was from Millipore Sigma (St. Louis, MO). NIH/3T3 cells were detached using trypsin and counted on a Countess II Automated Cell Counter (cat no. AMQAX1000, Thermo Fisher). Cells were grown and treated in a humidified 37°C, 5% CO_2_ incubator unless otherwise indicated.

### Transfection of *HMMR* mini-genes, RNA extraction, and cDNA synthesis

The day before the experiment, 125,000 NIH/3T3 cells/well were seeded into a 6-well plate (cat no. 08-772-1B, Thermo Fisher). Cells were seeded in antibiotic-free growth medium. The following day, cells were transfected with 500 ng *HMMR* reference minigene, *HMMR* sQTL alt minigene, or *HMMR* ancestral minigene plasmid DNA using TransIT-LT1 (cat no. 2304, Mirus Bio, Madison, WI) according to the manufacturer’s instructions; mock (no DNA)-transfected NIH/3T3 cells were used as the negative control. See plasmid section for complete sequence information. Cells were incubated for 48 h. After 48 h, cells were washed once with cold 1x PBS and harvested in 1 mL of cold 1x PBS by scraping. Cell suspensions were transferred to clean 1.5 mL tubes and pelleted by centrifugation (500 × g, 5 min, 4°C). Cell pellets were then washed once with 1 mL cold 1x PBS and pelleted by centrifuged (500 × g, 5 min, 4°C). After removal of supernatant, pellets were either processed immediately or stored at -80°C until needed. Five independent biological replicates were performed for each condition.

On the day of RNA extraction, cell pellets were thawed on ice and then immediately lysed using the QIAshredder kit (cat no. 79654, QIAGEN, Hilden, Germany). RNA was extracted from the flowthroughs using the RNeasy kit (cat no. 74104, QIAGEN), according to the manufacturer’s instructions. RNA was eluted in 30 µL RNase-free water, quantified on a NanoDrop 2000C (Thermo Fisher), and stored at -80°C until needed.

On the day of cDNA synthesis, RNA samples were thawed on ice. 200 ng of RNA was used as template for cDNA synthesis using the iScript Advanced cDNA Synthesis Kit (cat no. 1725038, Bio-Rad, Hercules, CA) according to the manufacturer’s instructions. A no RT reaction was prepared as a negative control. cDNA was diluted 1:2 in water and then used as a template for PCR or qPCR.

### PCR detection of *HMMR* mini-gene splice products

PCR was performed using the NEBNext High-Fidelity 2X PCR Master Mix (cat no. M0541L, New England Biolabs, Ipswich, MA) according to the manufacturer’s instructions with slight modifications. The total reaction volume was 25 µL, and 1 µL of diluted cDNA was used as template. PCR primers were validated for specificity in silico using Primer-BLAST (https://www.ncbi.nlm.nih.gov/tools/primer-blast/index.cgi), are listed in Table 1, and were from Integrated DNA Technologies (Coralville, IA). PCR reactions were combined with Novex Hi-Density TBE Sample Buffer (5X) (cat no. LC6678, Thermo Fisher) and then loaded onto a 20% Novex TBE gel (cat no. EC6315BOX, Thermo Fisher). The gel was stained with ethidium bromide (cat no. H5041, Promega Corporation, Madison, WI), washed several times with distilled water, and then bands were visualized on an E-gel Safe Imager (cat no. G6500, Thermo Fisher).

**Table 1.**
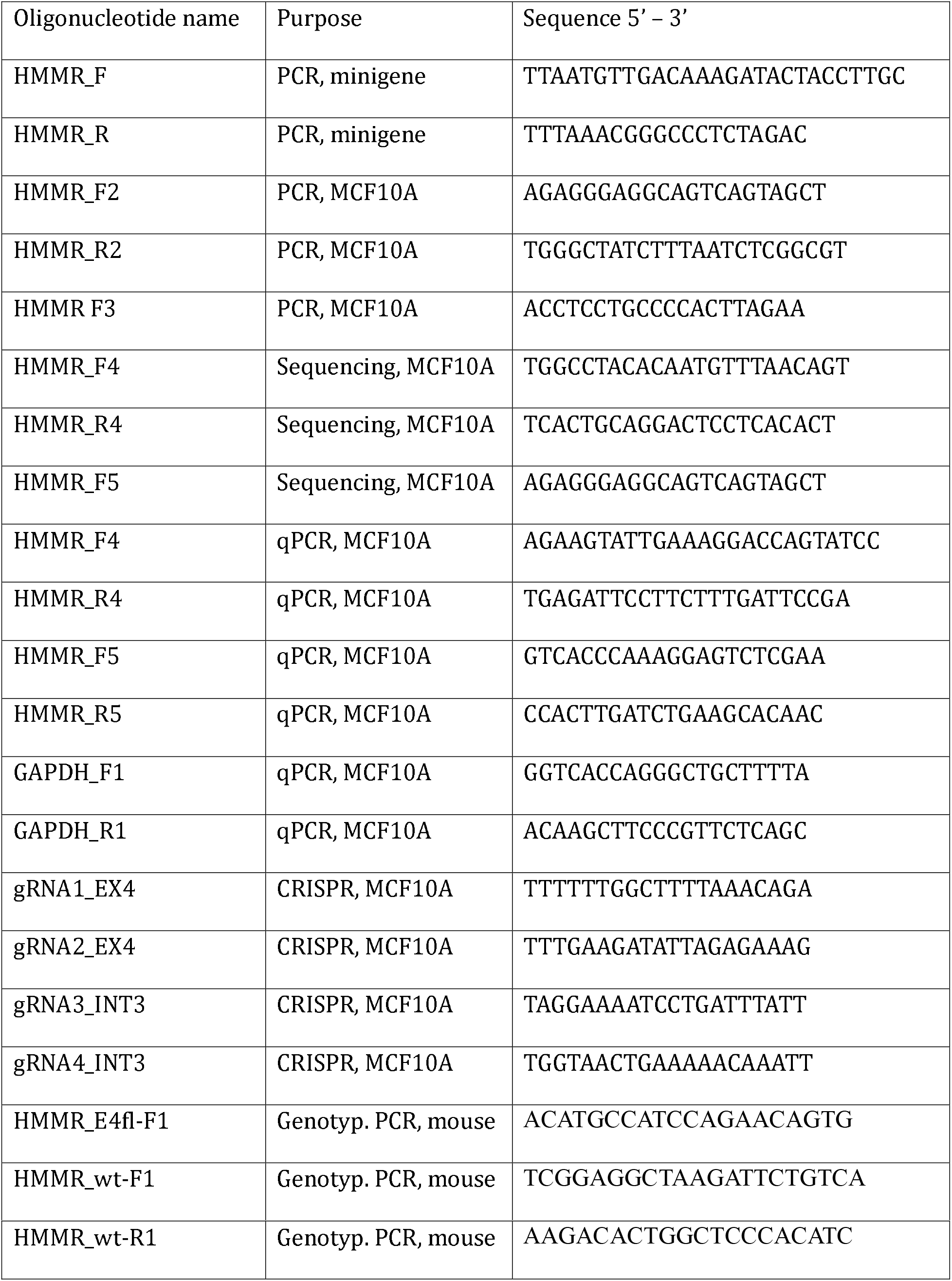
Oligonucleotide information.

Quantification of the splice junctions was done in ImageJ2 (version 2.16.0) as follows. For each lane, a fixed area region-of-interest (ROI) was drawn around the upper band (321 bp, which includes exon 4) and the integrated intensity was determined. This process was repeated for both the lower band (281 bp, which excludes exon 4) and a region below the lower band to determine the local background. Then the local background integrated intensity was subtracted from both the upper and lower band integrated intensities to generate corrected integrated intensities, which were used to calculate the exon 4-included:exon 4-excluded ratio. We log_10_ transformed these ratios and then use a paired t-test to test for a significant difference between genotypes.

### Cell culture

MCF10A RFP-TUBA1B cells were purchased from Sigma-Aldrich (CLL1039). MCF10A-RFPTUBA1B cells and CIRSPR edited clones were cultured in Brugge media (Debnath et al., 2003) comprised of DMEM/F12 (Gibco, 11320082) with 5% horse serum (Gibco, 16050122), 20 ng/mL human epidermal growth factor (EGF) (PreproTech, AF-100-15), 10 µg/mL bovine insulin (Sigma, I0516), 0.5 µg/mL hydrocortisone (StemCell, 7926), and 100 ng/mL cholera toxin (Sigma, C8052-2MG). Cells were incubated at 37°C and 5% CO2.

### Acini Culture

Cells were harvested from 70% confluent 6 well plates and live cells were counted using a haemocytometer and trypan blue dead cell stain. 1250 cells were resuspended in 40μL of Geltrex (Gibco, A1413202) and solidified into domes at 37°C for 15 minutes. The Geltrex domes were then incubated with Brugge media at 37°C and 5% CO2. Acini were cultured for 9 days and imaged using the EVOS M7000 Imaging system (Thermofisher). Acini area and circularity were quantified using ImageJ.

### Cas9 editing of MCF10A cells

Cas9 sgRNA were designed using IDT Alt-R CRISPR HDR Design Tool and ordered from IDT (Table 1). For each Cas9 reaction, 500,000 cells were resuspended in 100uL Opti-MEM (Gibco, 31985062), with 4uM ssODN HDR template and 2.4uM sgRNA-Cas9-GFP RNP complex. Cells were nucleofected with the RNP complex using the Amaxa Nucleofector II Device X-002 protocol. Nucleofected cells were resuspended in Brugge media and allowed to recover in the incubator. 24 hours post nucleofection, the BD FACS Fusion (Becton Dickinson) flow cytometer was used to perform single cell sorting of RFP+ GFP+ double positive cells into 96 well plates to isolate clones. Viable clones with the intended edit were identified via PCR genotyping (Table 1. Primers) and confirmed with sanger sequencing.

### Genotyping and sequencing

Cells were harvested from 70% confluent 6 well plates, and total gDNA was extracted with Qiagen Blood and Tissue (Qiagen, 69504). DNA concentration was quantified by the NanoDrop Microvolume Spectrophotometer. gDNA regions of interest were amplified via PCR (see Table 1 for primers) and amplicons were visualised with gel electrophoresis. PCR amplified gDNA regions were purified using QiaQuick PCR purification kit (Qiagen, 28104). Amplicons were sequenced with sanger sequencing (see Table 1 for primers) and sequencing reads were aligned to the HMMR sequence from GRCh38 (hg38) using Benchling.

### qPCR

Cells were harvested from 70% confluent 6 well plates, and total RNA was extracted with the RNeasy kit (Qiagen, 74104). RNA concentration was quantified by the NanoDrop Microvolume Spectrophotometer. Purified RNA was incubated with ezDNase enzyme to remove any gDNA contamination, and cDNA was then synthesized with SuperScript™ IV VILO™ Master Mix (Thermofisher 11756050). Samples reactions were prepared with Syber Green Master Mix (Applied Biosystems, 4364344), 0.25ng/μL cDNA, and 250nM of both the forward and reverse primers (see Table 1 for primers). qPCR was conducted using the QuantStudio™ 3 PCR System (Applied Biosystems) and fold changes were calculated with the 2^−ΔΔCt^ method, normalized to Gapdh as the housekeeping gene.

### Immunoblotting

Cells were harvested from 70% confluent 10 cm plates and lysed in RIPA buffer (Sigma R0278) supplemented with PhophoSTOP phosphatase inhibitor (Roche, 04906845001) and Protease Inhibitor Cocktail (Sigma, 4693159001). Protein concentration was determined using the Pierce™ BCA Protein Assay Kit (Thermofisher, 232225). Samples were separated by SDS-PAGE (4% stacking gel, 8% separating gel), then transferred to PVDF membrane. The membrane was blocked at room temperature with 3% BSA + PBST, then incubated with primary antibody (see Table 2 for antibodies) diluted in 3% BSA + PBST overnight at 4°C. The following day, the membrane was washed 3 times with PBST, then incubated with secondary antibody diluted in 3% BSA + PBST (see Table 2 for antibodies) for 2hr at room temperature. Membranes were imaged with Sapphire FL Biomolecular Imager (Azure Biosystems).

**Table 2.**
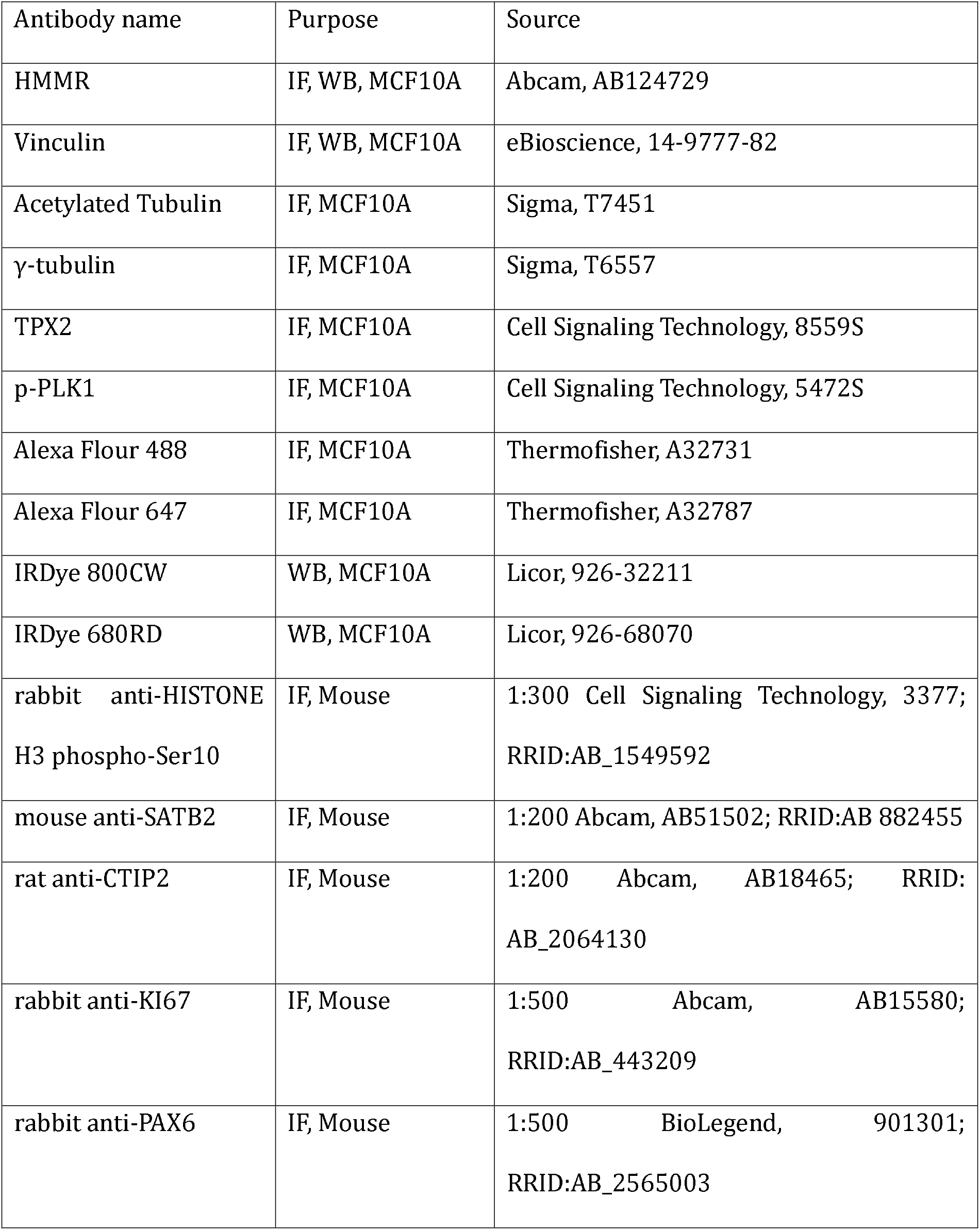
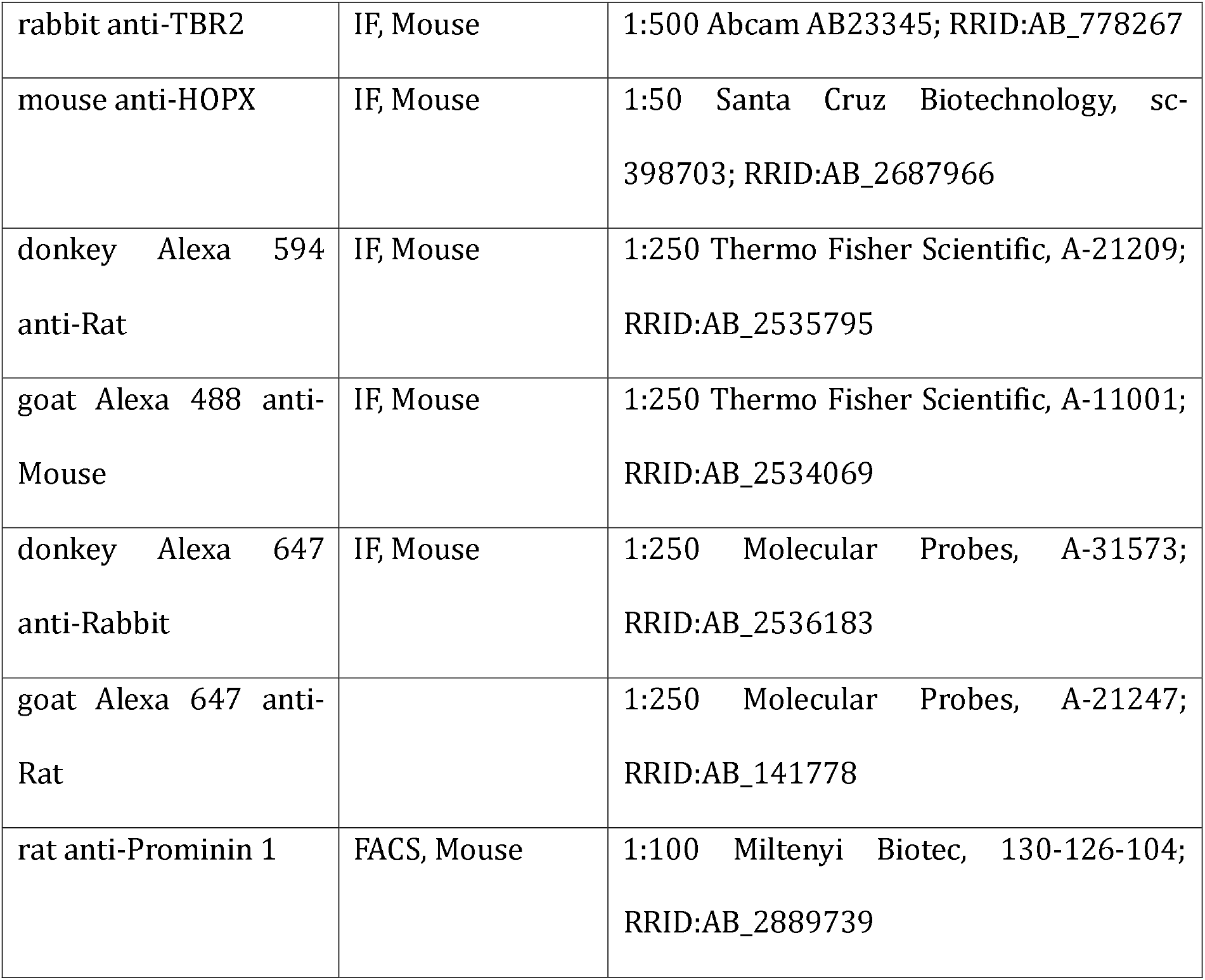
Antibody information.

### Colony forming assays

Cells were harvested from 70% confluent 6 well plates and live cells were counted using a haemocytometer and trypan blue dead cell stain. 100 cells were then plated into a single well in a 6 well plate and grown in Brugge media for 5 days at 37°C and 5% CO2. Each well was imaged on day 5 with the IncuCyte S3 (Satorius), 4x bright field whole well scan. Colony phenotype was determined by colony shape, cell morphology, and number of cell-cell contacts.

### Live cell imaging

Live cell imaging of day 3 colonies to assess cell division kinetics was performed with the EVOS M7000 Imaging system (Thermofisher) and EVOS Onstage Incubator. Day 3 colonies were imaged at 37°C and 5% CO2 with 80% humidity. Colonies were imaged every 3 minutes for 24 hours using the 20x objective, capturing both phase contrast and RFP channel wide-field images. Cell division phase was determined using the morphology of miotic spindle labelled with an endogenous RFP-tubulin tag.

### Immunofluorescence and confocal imaging for *in vitro* samples

Cells were seeded at 25,000 cells/well onto autoclaved glass coverslips in a 24 well plate and incubated for 48 hours until 70% confluent. Cells were washed twice with PBS and fixed for 3 minutes with 20°C methanol. Fixed cells on coverslips were blocked at room temperature with 3% BSA + PBST, then incubated with primary antibody (see Table 2 for antibodies) diluted in 3% BSA + PBST overnight at 4°C. The following day, coverslips were washed 3 times with PBST, then incubated with secondary antibody diluted in 3% BSA + PBST (see Table 2 for antibodies) for 2hr at room temperature. Coverslips were then washed 3 times with PBST, and incubated in 2μM Hoechst 33342 (Thermofisher, 62249) for 10 minutes at room temperature. Finally, coverslips were mounted onto slides with Prolong Gold Antifade Mounting Reagent (Life Technologies, P36935) and stored at 4°C.

Confocal imaging was captured with the Zeiss Cell Discoverer 7 Airyscan microscope. Slides were images using the 50x water-immersion objective and 1x tubulens. 10µm Z-stacks were captured at 0.5µm intervals, with both laser power and gain for each laser channel identical between samples. Fluorescence intensity was quantified using the ZEISS ZEN Microscopy Software.

### Generation of humanized *Hmmr* exon 4 skipping mouse model

Generation of genome-edited mice by Mouse Genetics Core Facility of Sloan Kettering Institute followed the protocol approved by the Institutional Animal Care and Use Committee (IACUC) of Memorial Sloan Kettering Cancer Center.

The conditional *Hmmr* allele was generated using Crispr/Cas9 with two gRNA and an ssDNA donor at the MSKCC transgenic core facility. Two LoxP elements (Left LoxP and right LoxP) are positioned 380 bp upstream and 327 bp downstream of exon 4. Mice were not bred to CRE lines so no LoxP recombination occurred in any of the analyzed mice. Mouse exon 4 and the 707 bp surrounding intronic sequences are replaced by the human sequence (partial Hmmr humanization). Mouse zygote donors are from C57BL/6J (Stock #000664; The Jackson Laboratories (JAX)) and in-house breeding thereof. Mouse surrogates for embryo transfer were in-house bred F1 females from C57BL/6 females (Stock #000664, #005304 or in-house bred offspring thereof) x CBA male (Stock #000656; JAX) crosses. Zygotes for microinjection of genome editing reagents were collected from superovulated C57BL/6 females copulated with C57BL/6 males. Those for electroporation were prepared by in vitro fertilization of oocytes from superovulated C57BL/6 females and sperm from C57BL/6 males.

Zygote pronuclear microinjection was performed to introduce genome editing cocktails (CRISPR mixes) containing a long single stranded DNA (lssDNA) knock-in (KI) donor. Zygotes were kept in a drop of M2 medium (Millipore-Sigma) during microinjection. CRISPR mixes for pronuclear injection consisted of Cas9 protein (50-100 ng/μl; IDT), crRNAs (#1 and #2, 25-50 ng/μl each; IDT), tracrRNA (50-100 ng/μl; IDT) and lssDNA donor (5-10 ng/μl; Megamer, IDT) in 1 mM Tris buffer (pH 7.4). The microinjection setup was composed of an inverted microscope (TE2000, Nikon), micromanipulators (TransferMan, Eppendorf), microinjectors (CellTram/FemtoJet, Eppendorf). In an attempt to specifically delete the interval between 2 gRNA target sites, electroporation was used for introduction of CRISPR mixes into zygotes without the lssDNA KI donor essentially as previously described (Takemoto et al., Dev Biol, 2016) with slight modifications. Briefly, zygotes prepared by IVF, 9-12 hours after insemination, were washed in electroporation buffer (0.01% poly vinyl alcohol (PVA) (Sigma-Aldrich) in Opti-MEM (Gibco-Invitrogen)) and placed in a glass slide-format platinum electrode (Protech International) filled with a CRISPR mix consisting of Cas9 protein (50-100 ng/μl; IDT), crRNAs (#1 and #2, 25-50 ng/μl each; IDT) and tracrRNA(50-100 ng/μl; IDT) in PVA/Opti-MEM electroporation buffer. The embryo-loaded electrode was attached to Genome Editor Electroporator (Protech International) applying 3 sets of reciprocal pulses (a total of 6 pulses) of 3 ms each with 97 ms intervals at 30V. Manipulated (microinjected or electroporated) zygotes were cultured in KSOM (Millipore-Sigma) until transfer into the oviduct of pseudopregnant surrogate females.

In total, two founder mice were backcrossed to wildtype mice and both lines were maintained independently. In the targeted allele of one founder, a 3-base nucleotide change (ACC to CAG) that is distinct from the lssDNA KI donor was detected at position -195-193 from the splice acceptor junction of human exon 4 (corresponding to chr5:163467490-163467492 in the hg38 reference genome). Throughout, we observed no major phenotypic differences between the two founder lines and always analyzed mice from both founder lines to ensure reproducibility of our analyses. The KI allele in the other founder was confirmed to be identical to the lssDNA KI donor and thus to the human reference genome.

The sequence of the lssDNA was (where upper case, underlined sequence is the human intronic sequence and the lower case sequence is the mouse intronic sequence acting as homology arms):

Gcttgccaacttaaactctgatgcaaataatcagtcgtagtctccgcagtgcgaccagcaatgtcactaagccatcactgcatgtg

Tgagcatcaaacatcagcctcactgccatggagcctcagaaaagaccagtgatactgaagtaaagagagaccggtgcctatg

ATAACTTCGTATAATGTATGCTATACGAAGTTAT<u>GCATACAGGAAGACTGTGAACGATG</u>

<u>GATAGATATAAATATTTGATCCACATATATAATTGTAATCAACTGACATAATGATTTTTCA</u>

<u>GTTATCAGTTTTTCATTAAGTGGGCTATCTTTAATCTCGGCGTTGCCAGCATACCACATT</u>

<u>AACATAGCCCGGATAGATCTGTGAAATACTGTAAGAACTGTTTTCCTGTAGGCACTGTT</u>

<u>CTGGATGGCATGTTTGCCTCACTGCAGGACTCCTCACACTGCAACAGAATCCTTATCTC</u>

<u>TTAAATCTGTGTAACTCTCTTTCTATCATGTGTACTTTTCACTAAGTTTAAAGCACTGCTT</u>

<u>AC</u>CTCTTTCTCTAATATCTTCAAATCTTTATCATTCTTTTGAGATTCCTT<u>CTGTTTAAAAG</u>

<u>CCAAAAAAACTTATAAGCAAATTTAGATGTCACAGATATGACAGCCAAATTTGTTTTTCA</u>

<u>GTTATCAGAAAAGATGAAGGGATCCAATCAGTTAGTTTCTATTTTAAACATTACACTGTT</u>

<u>AAACATTGTGTAGGCCACTCATCTGATAAATAAAATATTCTAAAAATCTCAAGACATTTTA</u>

<u>GGTTTAGAAATTTTCAACCCAATAAATCAGGATTTTCCTAATCAGCCCAGTGAAAAGAAC</u>

<u>AGGTGGAACTTTATGAATAAAGCTACTGACTGCCTCCCTCTAATGTTAAATAGCAACAC</u>

<u>ATCTTGAATATTTAATTATGCTATCTCAGAAAGAATCTTACAAAGAGAAATCGACTTAAAC</u>

<u>TTTGATACCC</u>ATAACTTCGTATAATGTATGCTATACGAAGTTATtgattagattcaaagaaaaattat

Aaccctacaacctatgatttttcatggcatttatttacataagacattcaatcttgtgtgtaaaacctttctgacacaatgaaagaaaat

gtagtttatacagtattaagaacaggtttgctgccaggcagtggtggcgcacgcctttaatcccagcactcgg

### Assessment of *Hmmr* exon 4 skipping in mouse model

To assess *Hmmr* exon 4 skipping, we grepped for chimeric kmers between mouse exon e and either mouse exon 4 (TGAGAGTCCTTCTTTGACACAG, kmer 1) or human exon 4 (TGAGATTCCTTCTTTGACACAG, kmer 2) as well as mouse exon 3 and mouse exon 5 (AGAGCGCGAATCTTTGACACAG, kmer 3). We then counted the number of reads supported by each kmer and computed the proportion of transcripts skipping exon 4 in *Hmmr*^+/+^ mice as the counts for kmer 3 divided by the sum of the counts for kmer 3 and kmer 1. For humanized mice, we used kmer 2 instead of kmer 1. This strategy avoids potential biases introduced by mapping to human or mouse exon 4.

### Mouse husbandry

All mouse procedures for neurobiology experiments were approved by the Austrian Federal Ministry of Science and Research in accordance with the Austrian and European Union animal law (license number: BMWF-66.018/0007-II/3b/2012, BMWFW-66.018/0006-WF/V/3b/2017, GZ 2024-0.698.056, and GZ 2025-0.597.515). Experimental animals were maintained and bred according to regulations approved by institutional animal care and use committee, institutional ethics committee and the guidelines of the preclinical facility at Institute of Science and Technology of Austria (ISTA). All mice used in this project showed specific pathogen free status according to FELASA recommendations^96^. The conditions were stablished as: 21±1°C ambient temperature and 40-55% humidity in 12 hrs dark/light cycles. Food (V1126, Ssniff Spezialdiäten GmbH, Soest, Germany) and tap water available ad libitum. Both male and female littermates of the desired genotypes were used randomly. Experimental mice were used at an age range from 2-8 months for breeding and at E14 and P21 for experiments. All efforts were made to minimize the number of animals used following the 3R principles. We have not observed any influence of sex on the results in our study. *Hmmr* mouse lines were kept in heterozygosity and in a C57/Bl6 genetic background at ISTA preclinical facility. Heterozygous mattings were placed in order to obtain the experimental samples (control and humanized mice). Siblings were used for the analysis.

### Mouse genotyping

Biopsies were collected from embryos and postnatal animals for genotyping. Genomic DNA was extracted using DirectPCR Lysis Reagent–Tail (Peqlab, Cat. No. 31-102-T). Samples were incubated overnight at 55°C with shaking at 300 rpm, followed by incubation at 85°C for 45 min. Genotyping PCR was performed using GoTaq® G2 DNA Polymerase Ready-to-Use Master Mix (Promega, Ref. M782B) according to the manufacturer’s instructions, with an annealing temperature of 60°C. PCR products were resolved and visualized by agarose gel electrophoresis. The *Hmmr* wild-type and humanized alleles were simultaneously detected using a three-primer PCR assay. HMMR_E4fl-F1, HMMR_wt-F1, and HMMR_wt-R1 primers (Table 1) were combined in the same reaction at a 1:1:1.5 ratio, respectively.

### Isolation of tissue and immunohistochemistry for mouse tissue

All antibody information is available in Table 2. Postnatal mice were deeply anesthetized by injecting a ketamine/xylazine solution (65 mg and 13 mg/kg body weight, respectively) and unresponsiveness was confirmed through pinching in the paw. The diaphragm of the mouse was opened from the abdominal side to expose the heart. Cardiac perfusion was performed with ice-cold PBS followed immediately by 4% PFA prepared in PB buffer (Sigma-Aldrich). Brains were removed and further fixed in 4% PFA o/n to ensure complete fixation. For mice embryos, mice pregnant females were sacrificed by cervical dislocation and embryonic mice heads were fixed by immersion in 4% PFA ice-cold for 3 hours. All brains were cryopreserved with 30% sucrose (Sigma-Aldrich) solution in PBS for approximately 48 hours. Brains were then embedded in Tissue-Tek O.C.T. (Sakura). For adult time points, 45µm coronal sections were collected in 24 multi-well dishes (Greiner Bio-one) and stored at -20°C in antifreeze solution (30% v/v ethyleneglycol, 30% v/v glycerol, 10% v/v 0.244M PO4 buffer) until used. Embryonic brains were sectioned at 20µm and directly mounted onto Superfrost glass-slides (Thermo Fisher Scientific) for storage at -20 C. For immunohistochemistry, adult brain sections were mounted, followed by 3 wash steps (5 minutes) with PBS. For EdU-KI67 and HOPX stainings, antigen retrieval protocol was performed by placing the sections in Citrate buffer 0.1M during 30 minutes at 75-85 °C and extra 30 minutes of cooling down. All tissue sections were blocked during one hour in a blocking buffer solution of 5% normal donkey serum (Thermo Fisher Scientific), 0.3% Trition X-100 in PBS and 2% BSA. Primary antibodies were prepared in blocking buffer and incubated o/n at 4°C. Sections were washed 3 times for 5 minutes each with PBS-T (0.3% Triton X-100 in PBS) and incubated with corresponding secondary antibody diluted in PBS-T for at least 2 hours. Sections were washed twice with PBS-T and once with PBS. Slices were incubated in 2.5% DAPI for nuclear staining (Thermo Fisher Scientific). Sections were embedded in mounting medium containing 1,4-diazabicyclooctane (DABCO; Roth) and Mowiol 4-88 (Roth) and stored at 4°C.

### Imaging and analysis of mouse tissue

Sections were imaged using either an inverted LSM880 or a Zeiss Slidescanner or Nikon CSU-W1-02 Spinning-disk and processed using Zeiss Zen Blue (RRID:SCR_013672) or ImageJ (RRID:SCR_003070) software. For image acquisition we used 20x objectives and z-stack for the entire thickness of the tissue. Tiled images were taken for at least three brain sections per animal and both hemispheres were quantified. Labeled cells were manually counted based on respective marker expression. Statistical analysis was performed in Graphpad Prism 11 (RRID:SCR_002798).

### Cortex area quantification

Cortical area was quantified from brightfield images of the whole brain using Fiji (ImageJ). For each sample, the cortical area of each hemisphere was manually delineated and measured. The areas of the left and right hemispheres were summed to obtain the total cortical area per brain. Measurements were performed for each genotype and used for subsequent statistical comparison.

### EdU labeling experiments

Cycling apical radial glial cells were assessed by EdU incorporation. Experiments were based on the use of the Click-iT Alexa Fluor 647 imaging kit (Thermo Fisher Scientific, Cat#C10340). Reagents were reconstituted according to the user manual. Intraperitoneal EdU injections were performed at E13, 24 hours before fixing (1mg/ml EdU stock solution; 30-40ml per mouse) to calculate the cell cycle exit. Tissue collection was followed by immunohistochemistry as described above.

### Spindle angle quantification in mouse cortex

Mitotic spindle orientation was quantified in PH3-labeled cortical sections using Fiji (ImageJ). PH3^+^ cells in late metaphase or early anaphase were identified, and the angle of the mitotic spindle relative to the ventricular surface was manually measured. For anaphase cells, the spindle axis was defined by the orientation of the two segregating chromosome masses. Mitotic cells located within 15 µm of the ventricular surface were classified as apical divisions, whereas cells located more than 15 µm from the ventricular surface were classified as basal divisions. Spindle orientation was categorized as horizontal (0–30°), oblique (30–60°), or vertical (60–90°). The distribution of division orientations was quantified and compared between genotypes.

### Preparation of single cell suspension for scRNA-Seq

Single cell suspension of cells prepared as described^97^. Experimental animals at E14 were sacrificed, brains extracted and cortex dissected. Each biological replicate corresponded to an individual animal. Control and humanized samples were obtained from littermates across four independent litters. Single cell suspensions were obtained by using Papain containing L-cysteine and EDTA (vial 2, Worthington, Cat#PAP2), DNase I (vial 3, Worthington, Cat#D2), Ovomucoid protease inhibitor (vial 4, Worthington, Cat#OI-BSA), EBSS (Thermo Fisher Scientific), DMEM/F12 (Thermo Fisher Scientific), FBS (Thermo Fisher Scientific) and HS (Thermo Fisher Scientific). All vials from Worthington kit were reconstituted according to the manufacturer’s instructions using EBSS. The dissected brain areas were directly placed into Papain-DNase solution (20units/ml papain and 1000 units DNase). Enzymatic digestion was performed for 15-30 minutes at 37°C in a shaking water bath. Next, solution 2 (EBSS containing 0.67mg Ovomucoid protease inhibitor and 166.7 U/ml DNase I) was added. The whole suspension was thoroughly mixed and centrifuged for 5 minutes at 1000rpm at RT. Supernatant was removed and cell pellet was resuspended in solution 2. Trituration with p1000 pipette was performed to mechanically disaggregate any remaining tissue parts. DMEM/F12 was added to the cell suspension as a washing solution, followed by a centrifugation step of 5 minutes with 1500rpm at RT. Cells were resuspended in media (DMEM/F12 containing 10% FBS and 10% HS) and kept on ice.

Immediately after generation of the cell suspension, cell concentration and viability were determined using an automated cell counter (Countess II/3 or Cellaca MX). Cells were fixed according to the Chromium Fixed RNA Profiling protocol (10x Genomics, CG000478, Rev. D). Cell pellets were resuspended in a solution containing 4% formaldehyde and 1× Fix and Perm Buffer and incubated at 4°C for 24 h. Following fixation, cells were pelleted, resuspended in 1× Quenching Buffer, and counted in an automated cell counter. Enhancer solution and glycerol were subsequently added, and samples were stored at −80°C until further processing.

Single-cell RNA-sequencing library preparation and sequencing were performed by the Next Generation Sequencing Facility at the Vienna BioCenter Core Facilities (VBCF-NGS), Vienna BioCenter (VBC), Austria. Libraries were prepared using the 10x Genomics Chromium Fixed RNA Profiling (10X Genomics Flex v1 workflow) workflow according to the manufacturer’s instructions (CG000527). Libraries were sequenced on an Illumina NovaSeqX platform.

### Preparation of single cell suspension for bulk RNA-Seq

Experimental animals were sacrificed at E14, and the brains were extracted and cortices dissected. Each biological replicate corresponded to an individual animal. Control and humanized samples were obtained from littermates across four independent litters. Cortical tissue was dissociated into single-cell suspensions using the Neural Tissue Dissociation Kit (P) (Miltenyi Biotec, #130-092-628), with stock reagents prepared according to the manufacturer’s instructions. Briefly, dissected cortices were collected in ice-cold Ca²⁺/Mg²⁺-free HBSS and centrifuged at 300g for 2 minutes at room temperature. Tissue was resuspended in pre-activated Enzyme Mix T and incubated for 15 minutes at 37°C with continuous shaking at 300 rpm. Samples were then mechanically dissociated by gently pipetting up to 10 times with a wide-tipped, fire-polished Pasteur pipette. Subsequently, pre-activated Enzyme Mix A was added, and samples were incubated for an additional 10 minutes at 37°C and 300rpm. Tissue was further dissociated by gently pipetting up to 10 times using a fire-polished Pasteur pipette with a reduced tip diameter, while avoiding the generation of air bubbles. The resulting cell suspension was passed through a 70µm cell strainer and washed with 10 mL Neurobasal Medium (NBM; Gibco). Cells were centrifuged at 300g for 10 minutes at room temperature. The supernatant was carefully removed, leaving approximately 1–2 mL to avoid disturbing the cell pellet. Cells were gently resuspended and washed in NBM, followed by centrifugation at 300g for 10 minutes at room temperature.

For enrichment of cortical progenitor cells, single-cell suspensions were stained with PE-Vio770-conjugated anti-Prominin-1 antibody (1:100) in a final staining volume of 200 µL. Appropriate unstained and single-stained controls were prepared in parallel. Samples were incubated for 30–40 minutes on ice with gentle mixing every 5–10 minutes. Cells were subsequently washed with 3 mL NBM and centrifuged at 300g for 10 minutes at room temperature. Prior to fluorescence-activated cell sorting (FACS), stained cells were resuspended in 500 µL NBM supplemented with serum [approximately 8.9% (v/v) horse serum and 8.9% (v/v) heat-inactivated fetal bovine serum], while control samples were resuspended in 200 µL of the same medium. Right before sorting, cell suspension was filtered using a 40mm cell strainer. Samples were then immediately subjected to FACS for isolation of Prominin-1^+^ single cortical progenitor cells for downstream bulk RNA-seq. FACS was performed on a Sony SH800SFP (SH800S software, RRID:SCR_018066) using 100 nozzle and keeping sample and collection devices (5ml tubes) at 4°C. Duplet exclusion was performed to ensure sorting of true single cells. After sorting, samples were centrifuge at 300g during 10 minutes. Supernatant was removed and RNA extraction was performed as described^97^ using custom made lysis buffer and Trizol but omitting the use of GlycoBlue and the over night precipitation step. RNA quality was analyzed using Bioanalyzer RNA 6000 Pico kit (Agilent) following the manufacturer’s instructions. For library generation, 200 ng of high-quality total RNA was used as input for the Watchmaker mRNA Library Prep Kit (Watchmaker Genomics, Cat. #7BK0001). Libraries were prepared at the Vienna BioCenter Core Facilities (VBCF Vienna) following the protocol provided by the manufacturer. The resulting libraries were sequenced on an Illumina NextSeq 2000 platform by the Next Generation Sequencing Facility at VBCF.

### Analysis of bulk RNA-sequencing data

We aligned bulk RNA-seq data to the GRCm39 reference genome using STAR v2.7.8a^78^ with parameters --quantMode TranscriptomeSAM --quantTranscriptomeBan IndelSoftclipSingleend --outSAMattributes MD NH --outSAMtype BAM Unsorted --runThreadN 8 --readFilesCommand zcat --outFilterMultimapNmax 1. We then used samtools^98^ v1.9 to sort and index the resulting file and umi_tools^99^ to deduplicate reads. We then used the RSEM^100^ v1.3.3 function rsem-calculate-expression with parameter –paired-end to quantify transcript expression. We excluded data from one sample as it was a clear gene expression outlier relative to the other seven samples (mean Pearson’s rho with other samples of 0.94 compared to > 0.98 for all other samples). We aggregated transcript level counts to gene level with tximport^101^ v1.36.1 and then tested for differential expression with DESeq2^102^ v1.48.1, including mouse sex (inferred from the expression of Y-linked genes) as a binary covariate. During this initial testing, we noticed that several ganglionic eminence marker genes had systematically higher expression in the samples from humanized mice. Although this could reflect genuine biological differences between the mice, we were primarily interested in differences in glutamatergic neuron progenitors. Therefore, we generated a ganglionic eminence (GE) expression signature score based on established GE marker genes (*Dlx1*, *Dlx2*, *Dlx5*, *Nkx2-1*, *Lhx6*, and *Nr2f2*). Expression values were variance-stabilizing transformed using DESeq2, and the GE signature score for each sample was calculated as the mean expression of these genes after transformation, followed by z-score normalization across samples. This score was included as a continuous covariate in the DESeq2 design formula to account for variation in GE-associated transcriptional programs.

### Analysis of single cell RNA-sequencing data

Initial analysis of the scRNA-seq data proceeded with the standard scanpy^103^ v1.11.4 workflow. Briefly, this involved filtering out cells with greater than 15% mitochondrial counts or n_genes_by_counts < 7500, log normalizing the data, identifying highly variable genes (with parameters min_mean=0.0125, max_mean=3, min_disp=0.5), running principal components analysis (PCA), identifying nearest neighbor cells (with parameters n_neighbors=10, n_pcs=40), and then performed Leiden^104^ clustering at resolution 0.4. Gross cell types were classified based on the expression of the following marker genes: GABAergic neurons as *Gad2*+^105^, glutamatergic neurons as *Slc17a6*+ or *Slc17a7*+^106^, progenitors as *Nes*-high and *Vim*-high^107^, Cajal-Retzius cells as *Reln*+^108^, and vascular cells as *Pecam1*+^109^ or *Pdgfrb*+^110^. Within progenitors, radial glia were identified as *Pax6*-high^51^, intermediate progenitors as *Eomes*+^51^, and GABAergic (i.e. ganglionic eminence) progenitors as *Dlx2*+^111^. Inhibitory neurons were classified according to their ganglionic eminence (GE) of origin with medial GE neurons being *Erbb4*+^112^ and lateral GE neurons being *Isl1*+^113^. We were unable to cleanly identify caudal GE neurons.

From there, we subset to only the two clusters of radial glia. We then reclustered the data with leiden resolution = 0.8 and removed small, clear doublet clusters that were double positive for *Pax6* and one of the above markers, removed all *Eomes*+ cells, and removed all *Nr2f2*+ cells. We then pseudobulked counts per sample by summing the raw counts for all cells passing the above filtering criteria in each sample and then tested for differential expression using DESeq2 and the above pipeline, without regressing out the GABAergic signature. We then removed all genes that were expressed in less than 10% of cells. To test for differences in the number of *Hopx*-high cells, we further required non-zero *Pax6* expression and then computed the proportion of cells in each sample with log normalized *Hopx* counts greater than 2, pairing samples by their founder of origin for the t-test.

### Combined analysis of bulk and single cell RNA-sequencing data

We joined the bulk and radial scRNA-seq DESeq2 outputs. Throughout, we used the sign of the log_2_ fold-change multiplied by the -log_10_(p-value) as a measure of both the direction of the change in expression and the confidence in it. We restricted further analysis to genes tested in both datasets that had the same sign for this statistic. We used a previously published set of genes with enriched expression in outer radial glia relative to other radial glia^55,58^ in testing for systematically higher expression of outer radial glia marker genes in the humanized mice.

### Statistics

For statistics related to analysis of RNA-seq data (i.e. Fig. 1-2, and part of Fig. 5), the implementation in scipy v1.15.3 was used. For *in vitro* and *in vivo* experiments, GraphPad prism was used to conduct all statistical tests. One-way-ANOVA tests were performed to compare the means of more than two groups. Chi-squared tests were performed to determine significance between the categorical variables planar vs. non-planar cell divisions or luminal vs. basal colony phenotypes and the spindle distribution in mouse tissue. Unpaired t-test was performed to compare between control and humanized mice when quantifying for cortical area, cortical thickness, cell-cycle exit, PH3, PAX6, TBR2, HOPX and spindle angle distribution. Two-way ANOVA was performed to assess significance for SATB2/CTIP2 quantifications.

## Acknowledgements

We would like to acknowledge the MSKCC transgenic mouse core facility for generating humanized mice, as well as Nancy Du, Xiang Chen, and other Du lab members for mouse breeding and genotyping. This research was supported by the Scientific Service Units (SSU) of IST Austria through resources provided by the Imaging and Optics- (IOF), Lab Support- (LSF) and Preclinical Facilities (PCF). We would also like to thank Fee Wielath and Kerstin Feistel for helpful discussions. Some figures were made with biorender.

## Data availability

Bulk and single cell RNA-sequencing data generated in this study have been uploaded to the Gene expression omnibus (GEO) with accessions GSE346330 and GSE346338 respectively.

Bulk RNA-seq data used to identify htCONDELs are publicly available with the following GEO accessions: GSE146481, GSE144825, and GSE232949. Splicing data from additional species was downloaded from https://apps.kaessmannlab.org/alternative-splicing/ and GTEx v10 sQTL data was downloaded from https://gtexportal.org/api/v2/association/dynsqtl.

## Code availability

Code for this study is available at: https://github.com/astarr97/Splicing. Code to perform the alignment of RNA-seq data from hybrid cells is available at: https://github.com/banwang27/multi-celltypes.

## Author contributions

A.L.S performed all analysis related to the identification of (i)htCONDELs. J.R. performed all experiments with MCF10A cells. A.V., Y.C., and F.M.P. performed all analysis related to the mouse model with the exception of the analysis of single cell and bulk RNA-sequencing data which was performed by A.L.S. A.L.S., A.V., J.R., C.M., S.H., and H.B.F. conceived the study, wrote the manuscript, and made figures. All authors approved the final version of the manuscript.

## Description of supplemental files

Supplemental_Figures.docx: Contains all supplemental figures.

Table_S1.csv: Contains information on transcriptomic deletions identified in this study.

Table_S2.csv: Contains information on indirect human transcriptomic deletions identified in this study.

