## Supplemental_Figures for "A human transcriptomic deletion links cortical expansion and cancer"

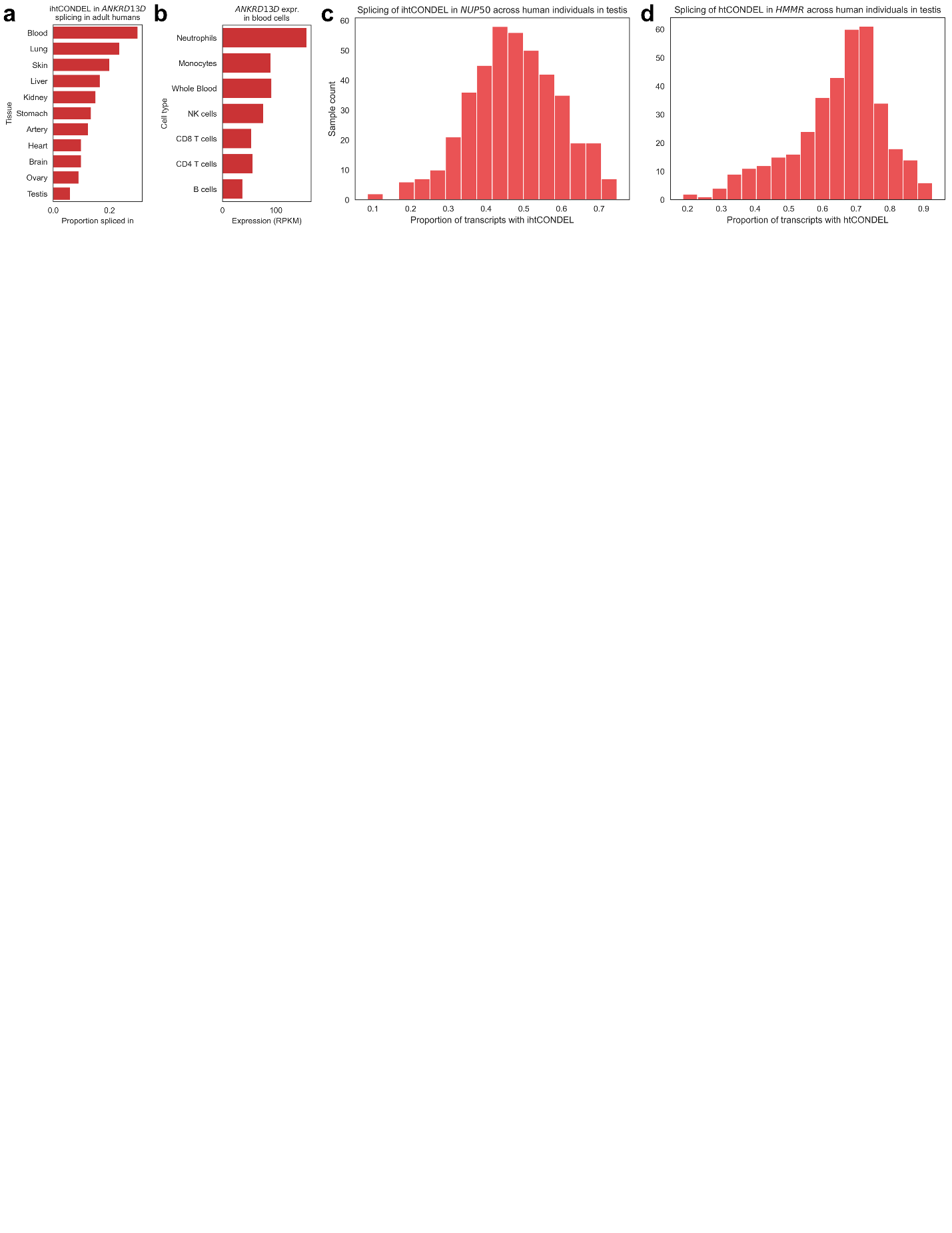

**Figure S1 (related to figures 1 and 2): a)** Proportion spliced in for ihtCONDEL-causing exon inclusion tissues for *ANKRD13D*. **b)** *ANKRD13D* expression in blood cell types. **c)** Proportion of transcripts with the ihtCONDEL in *NUP50* in testes across GTEx individuals. **d)** Proportion of transcripts with the htCONDEL in *HMMR* in testes across GTEx individuals.

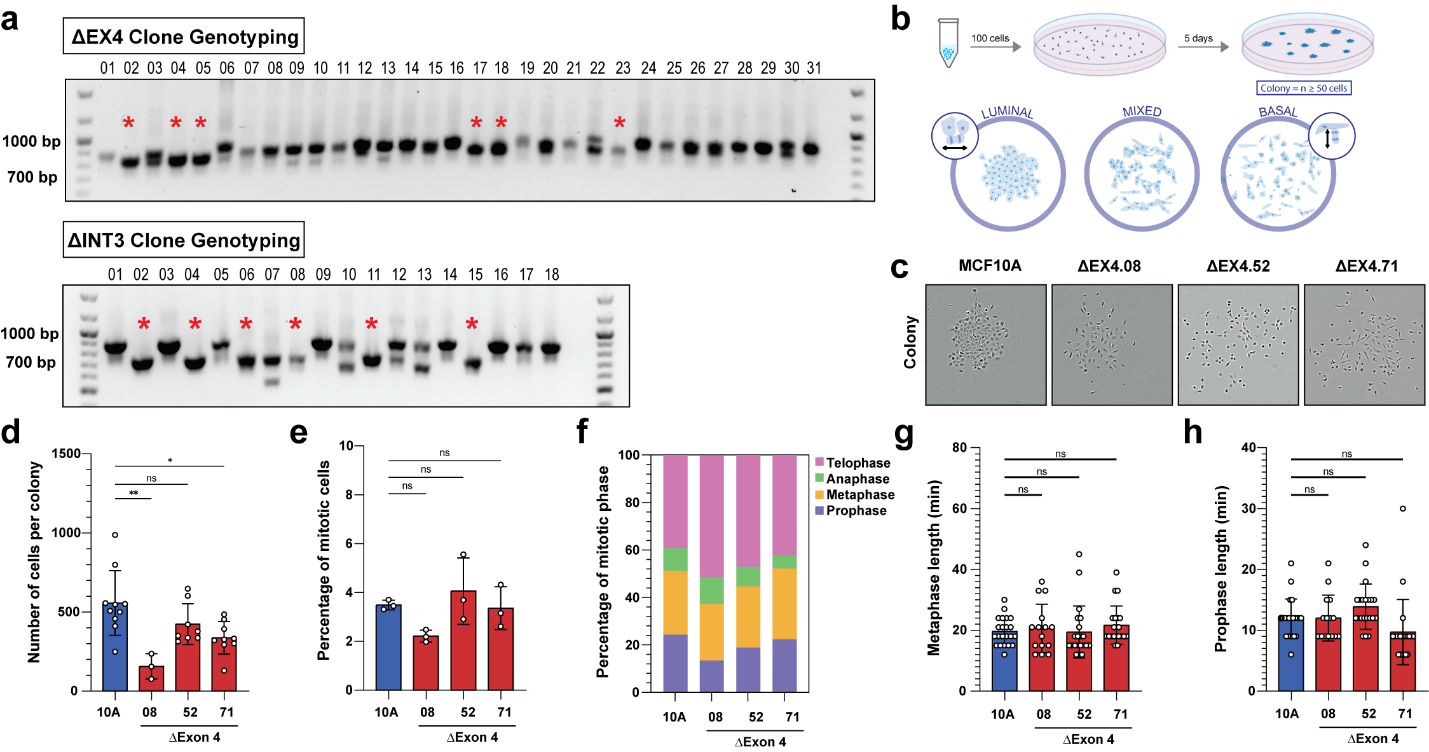

**Figure S2 (related to figure 3):** **a)** PCR genotyping of single cell clones screened for ΔEX4 and ΔINT3 CRISPR Cas9 edits. Red stars indicate clones selected for sanger sequencing analysis. **b)** Diagram showing the three colony phenotypes (luminal, mixed, and basal) with a flow chart of the colony forming assay workflow above. Luminal colonies are composed of tightly packed cuboid cells with many cell-cell contacts. Basal colonies have more elongated spindly cells, with fewer cell-cell contacts. Mixed colonies have both luminal and basal phenotypes. **c)** Day 5 colony forming assays with representative images of individual colonies, and **d)** number of cells per colony (one-way ANOVA). **e)** Percentage of mitotic cells (One-way ANOVA, n=3 replicates) and **f)** percentage of mitotic phase (One-way ANOVA), n=3 replicates). **g)** Length of metaphase, and **h)** length of prophase (One-way ANOVA, n=20 cells per genotype) from live cell day 3 MCF10A RFP-TUBA1B colonies. For all plots, statistical tests are indicated in the legend. Statistical significance is represented as follows ** = p<0.01, * = p<0.05, ns = p>0.05.

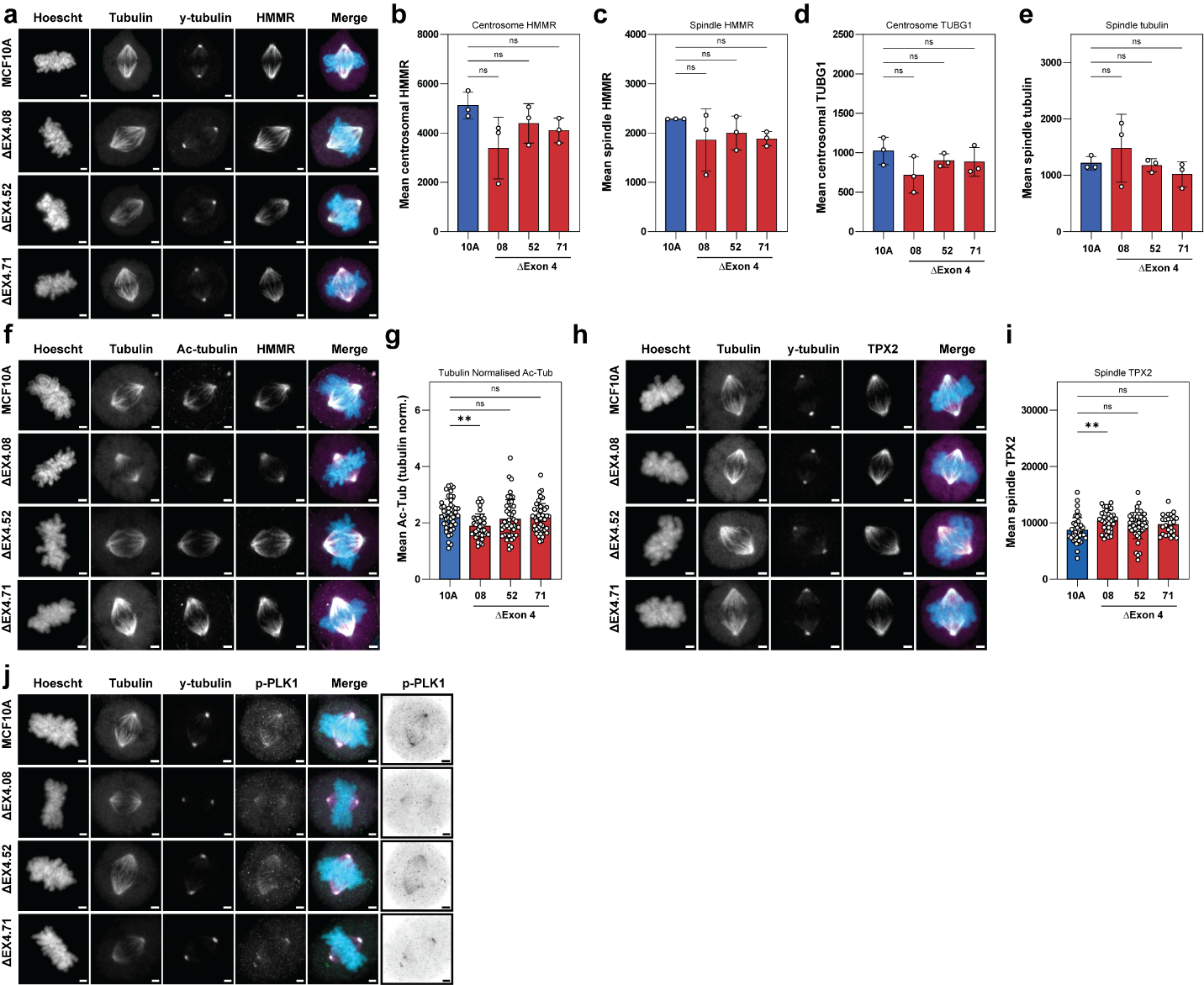

**Figure S3 (related to figure 3):** **a)** Confocal immunofluorescence images of metaphase cells stained for DNA (Hoechst), y-tubulin (TUBG1), tubulin, and HMMR, with a merged image of all channels. **b)** Quantification of mean HMMR fluorescence at the metaphase spindle poles, and **c)** the metaphase spindle (One-way ANOVA, n=3 replicates, n=30 cells per replicate). **d)** Quantification of mean TUBG1 fluorescence at the metaphase spindle poles, and **e)** mean tubulin fluorescence at the metaphase spindle (One-way ANOVA, n=3 replicates, n=30 cells per genotype). **f)** Confocal immunofluorescence images of metaphase cells stained for DNA (Hoechst), y-tubulin (TUBG1), tubulin, and Ac-tubulin, with a merged image of all channels. **g)** Quantification of mean Ac-tubulin fluorescence at the metaphase spindle (One-way ANOVA, n=30 cells per genotype). **h)** Confocal immunofluorescence images of metaphase cells stained for DNA (Hoechst), y-tubulin (TUBG1), tubulin, and TPX2, with a merged image of all channels. **i)** Quantification of mean TPX2 fluorescence at the metaphase spindle (One-way ANOVA, n=30 cells per genotype). **j)** Confocal immunofluorescence images of metaphase cells stained for DNA (Hoechst), y-tubulin (TUBG1), tubulin, and phosphorylated PLK1 (p-PLK1), with a merged image of all channels. For all plots, statistical tests are indicated in the legend. Statistical significance is represented as follows **** = p<0.0001, *** = p<0.001 ** = p<0.01, * = p<0.05, ns = p>0.05.

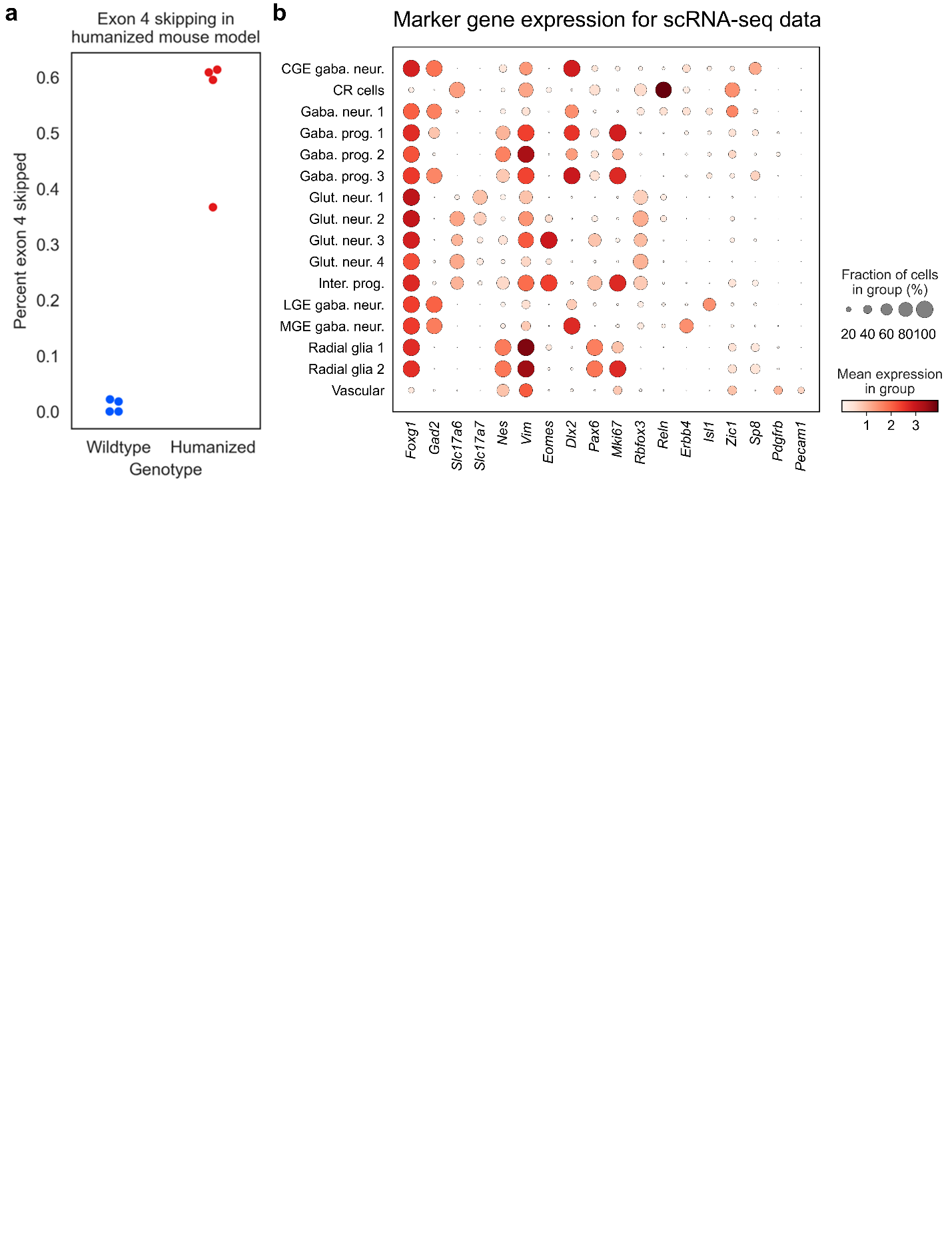

**Figure S4 (related to figure 5): a)** Rate of exon 4 skipping in wildtype (right, blue) and humanized (left, red) embryonic mouse forebrain progenitor cells. **b)** Expression of marker genes used to identify cell types in scRNA-seq data. C/L/MGE = caudal/lateral/medial ganglionic eminence, CR = Cajal-Retzius, neur. = neurons, prog. = progenitors, Glut. = glutamatergic, Inter. = intermediate.
